# Transmitted α-synuclein extracellular vesicles downregulate axonal flux of retrograde carriers in recipient neurons

**DOI:** 10.1101/2025.09.22.677685

**Authors:** Barbara E Duda, Saber H. Saber, Vanessa Lanoue, Adekunle T. Bademosi, Rachel S Gormal, Tristan P. Wallis, Alex J. McCann, Yara Abosnwber, Justin J. Cooper-White, Patrik Verstreken, Frédéric A. Meunier

**Affiliations:** Clem Jones Centre for Ageing Dementia Research (CJCADR), Queensland Brain Institute (QBI), The University of Queensland, St Lucia Campus, Brisbane, QLD 4072, Australia; Molecular Cell Biology Lab, Zoology and Entomology Department, Faculty of Science, Assiut University, Assiut 171516, Egypt; Tissue Engineering and Microfluidics Laboratory, Australian Institute for Bioengineering and Nanotechnology, University of Queensland, St Lucia Campus, Brisbane, QLD 4072, Australia; School of Chemical Engineering, University of Queensland, St Lucia Campus, Brisbane, QLD 4072, Australia; VIB Centre for Brain & Disease Research, KU Leuven, 3000 Leuven, Belgium; Department of Neurosciences, Leuven Brain Institute, KU Leuven, 3000 Leuven, Belgium; School of Biomedical Sciences, The University of Queensland, St Lucia Campus, Brisbane, QLD 4072, Australia

**Keywords:** α-Synuclein, Conditioned Medium, Axonal Retrograde Transport, Parkinson’s Disease, Exosome, Super Resolution Microscopy

## Abstract

α-Synuclein (α-syn) is a cytosolic protein located in nerve terminals and is involved in several neurodegenerative diseases such as Parkinson’s disease. Recent studies have demonstrated that α-syn can be transmitted from neuron to neuron via exosomal release, thereby contributing to the propagation of α-syn pathology. However, the mechanism by which α-syn-containing exosomes perturb the function of recipient neurons is currently unknown. Retrograde axonal transport of carriers emanating from the presynapse is essential for neuronal survival. To determine the effect of transmitted α-syn on neuronal retrograde trafficking, we used conditioned medium from α-syn transfected “donor” hippocampal neurons, applied to naïve (non-transfected) “recipient” neurons cultured in microfluidic chambers. Time-lapse imaging of retrograde carriers containing cholera toxin-B subunit (CTB) was then performed in these recipient neurons. Here, we show that conditioned medium from α-syn^WT^-GFP transfected donor neurons significantly downregulated the frequency of retrograde CTB carriers when applied to recipient neurons. This effect was abolished by (i) inhibiting endocytosis in recipient neurons, (ii) by blocking exosomal release from donor cells via an Hsp90-dependent mechanism, or (iii) by using conditioned medium from neurons transfected with a Parkinson’s mutant (α-syn^A30P^-GFP). Whilst transmitted α-syn-mEos2, α-syn^A30P^-mEos2 and mEos2 alone were detected in recipient neurons using single molecule imaging, interestingly, α-syn^WT^-mEos2 exhibited lower mobility, and periodic (190 nm) immobilisations along the axon. Our data suggest that the downregulation of vesicular trafficking by transmitted α-syn does not rely on the specific release nor uptake of exosomal α-syn but probably depends on its interaction with endogenous α-syn following endocytosis in recipient neurons. The transmitted α-syn containing extracellular vesicles therefore controls essential axonal trafficking in recipient neurons, an effect lost with the Parkinson’s disease mutant α-syn^A30P^.

## Introduction

The ability of neurons to survive, some of them for an organism’s entire lifetime, is a phenomenon that has been intensely studied. A neuron’s decision to either survive or degenerate relies on its ability to interpret signals from its environment. Neurons accomplish this feat through unique ligand receptor signalling, endocytosis of external factors, by integrating neuronal communication inputs from their connected network, along with surrounding cues. Critical to this is the ability of distally internalised factors to inform the cell body via long range axonal retrograde transport (Wang et al. 2015; Wang et al. 2016; Maday and Holzbaur, 2012; Ye et al. 2003). Many neurodegenerative diseases are characterised by the spread of toxic protein species, suggesting a role for neuronal transport in this process. Indeed, Braak et al. hypothesised that the propagation of Lewy bodies, a neuropathological hallmark of Parkinson’s disease (PD), multiple systems atrophy and Lewy body dementia, might be correlated with the progression of disease (Braak et al. 2003). Lewy bodies result from the accumulation of fibrillary forms of alpha-synuclein (α-syn) inside cytoplasmic inclusions (Spillantini et al., 1998; Spillantini, 1999). The spreading hypothesis gained momentum a few years later following the identification of a host-to-graft transmission of α-syn (Li et al. 2008). This study revealed, in deceased PD patients, the presence of Lewy body-like inclusions in transplants that were grafted a decade earlier. This ability of α-syn to spread was then confirmed *in vitro* (Hansen et al. 2011; Reyes et al. 2015) and *in vivo* (Angot et al. 2012; Luk et al., 2012; Masuda-Suzukake et al. 2013; Peelaerts et al. 2015, Okuzumi et al. 2018). Furthermore, brain homogenates from multiple system atrophy, dementia with Lewy bodies or Parkinson’s disease patients were shown to induce neurodegeneration when injected in the brain of mice or monkeys (Watts et al. 2013; Prusiner et al. 2015; Ayers et al. 2022; Bourdenx et al. 2020). Demonstration of α-syn transmissibility, its ability to recruit endogenous α-syn as well as to induce pathology in cells and mammalians exposed to abnormal α-syn, suggested a prion-like property of α-syn.

α-Syn is a soluble cytoplasmic protein consisting of 140 amino-acids with a molecular weight of 14 kDa, which is enriched at the presynapse (Maroteaux et al. 1988; Iwai et al. 1995; Yang et al. 2010). It exists in multiple forms from monomeric (Weinreb et al. 1996; Conway, Harper, and Lansbury 1998; Fauvet et al. 2012) through to complex multimeric structures (Burré et al. 2013; Wang et al. 2014). Several mechanisms have been implicated in the transmission of α-syn (Vasili, Dominguez-Meijide, and Outeiro 2019; Valdinocci et al. 2017; Hijaz and Volpicelli-Daley 2020). Exosomal release has emerged as a potential pathway in the spreading of α-syn for synucleopathies, as well as other cytosolic amyloid proteins involved in neurological disorders including Alzheimer’s disease (Howitt and Hill 2016; Liu et al. 2019). Exosomes are extracellular vesicles that play an important role in intercellular communication by transporting a variety of cytosolic molecules (Bobrie et al. 2012; van Niel, D’Angelo, and Raposo 2018; Gurung et al. 2021). They are formed following the invagination and fission of intralumenal vesicles from the endosomal membrane thereby generating multivesicular bodies (MVB). Upon fusion of an MVB with the plasma membrane, the intralumenal vesicles are released as exosomes (Théry, Zitvogel, and Amigorena 2002; Gurung et al. 2021). Evidence that α-syn could be transmitted via exosomes was first shown in the SH-SY5Y neuroblastoma cell line (Emmanouilidou et al. 2010), where conditioned medium from cells expressing α-syn increased cell death when applied to recipient cells. Moreover, exosomes isolated from the conditioned medium of primary cortical neurons were shown to contain oligomers of α-syn and to induce more apoptosis when added to H4 neuroglioma cells compared to free oligomers (Danzer et al. 2012). The ability of exosomes to transmit α-syn pathology was confirmed *in vivo* with the injection of exosomes extracted from the brain of Lewy body dementia patients into the brain of wild-type (WT) mice induced α-syn pathology (Ngolab et al. 2017). Furthermore, studies on exosomal α-syn in PD patients revealed different levels of α-syn in body fluids compared to healthy patients and the level of exosomal α-syn correlated with disease progression (Shi et al. 2014; Stuendl et al. 2016; Cerri et al. 2018; Jiang et al. 2020). Taken together, these studies suggest that exosomal release of α-syn from donor neurons is likely to contribute to the pathomechanism of neurodegeneration in PD. However, the mechanism of α-syn exosomal release from donor neurons, and how exosomes deliver their content into recipient neurons is not fully understood. Further, the impact of uptaken α-syn in recipient neurons and its long-term effect on survival remains to be established.

We previously demonstrated that presynaptic activity controls the retrograde trafficking of autophagosomes (Wang et al. 2015) and signalling endosomes (Wang et al. 2016; Wang et al. 2020). In this study, we investigated the impact of exosomally-transmitted α-syn on the long-range retrograde transport in recipient neurons. Using a dynamin inhibitor, we also tested the dynamin-dependency of the uptake of exosomal α-syn in recipient neurons. We also tested whether α-syn exosomal release was dependent on the activity of the heat shock protein 90 (Hsp90) in donor hippocampal neurons, which has recently been shown to be implicated in the fusion of MVBs to the plasma membrane in a *Drosophila melanogaster* model (Lauwers et al. 2018). Finally, we used single molecule super-resolution microscopy to track transmitted α-syn in recipient neurons and established its nanoscale organisation. These studies were performed using α-syn wildtype (α-syn^WT^) and α-syn^A30P^, a familial mutation of α-syn associated with PD.

## Results

To examine the impact of transmitted α-syn on synaptic trafficking, we expressed α-syn-GFP in hippocampal neurons (DIV12), harvested their media after 24h and applied it on untransfected (naïve) hippocampal neurons (DIV13) cultured in microfluidic chamber (Taylor et al. 2005). To test whether such conditioned medium could affect presynaptic trafficking and axonal transport involved in neuronal survival, we used Cholera toxin subunit B (CTB) conjugated to Alexa Fluor™ 555 (CTB-Af555), which was previously shown to be uptaken at the presynapse and transported via long range retrograde axonal transport to the cell body within signalling endosomes (T. Wang et al. 2016).

The conditioned medium from the soma chamber was applied to the nerve terminal chamber of the microfluidic device containing naïve neurons with a 70:30 vol/vol ratio (Figure 1 A). The conditioned medium was removed after 24 hours and CTB-Af555 (50 ng/mL) was added for 5 minutes to the nerve terminal chamber in high potassium (HK^+^) buffer to stimulate uptake of CTB in nerve terminals (T. Wang et al. 2015). Chambers were then washed three times in low potassium (LK^+^) buffer, and the conditioned medium was added back for another 2 hours at 37°C. The number of CTB-positive (CTB+) carriers undergoing fast retrograde axonal transport was examined by time-lapse confocal imaging of axons located in microgroove channels. In recipient (untransfected) neurons exposed to conditioned medium from α-syn^WT^-GFP-expressing neurons, we observed a significant decrease in the number of CTB+ carriers undergoing retrograde transport per second compared with neurons exposed to conditioned medium from neurons expressing EGFP (Figure 1 B and C). This suggests that the conditioned medium from α-syn^WT^-GFP-expressing neurons can negatively regulate the frequency of signalling endosomes that are transported from the presynapse to the cell body carrying the survival signal. This regulation is likely capable of controlling the number of signalling endosomes that are transported to the cell body. Indeed, the percentage of colocalization of CTB with TrkB-positive carriers was largely increased in axons preincubated with conditioned medium from neurons expressing α-syn^WT^-GFP compared to conditioned medium from neurons expressing EGFP (Supp Fig. 1 A). No difference was observed between conditions for the colocalization of CTB with LC3-positive carriers (Supp Fig. 1 B). We next examined whether PD human mutation A30P, had similar effect on CTB retrograde transport. Interestingly, the conditioned medium from α-syn^A30P^-GFP expressing neurons did not significantly impact CTB-Af555 retrograde trafficking, suggesting that the PD-linked mutation prevented this regulatory mechanism from occurring. Interestingly, in those axons the percentage of colocalization between CTB with TrkB-positive carriers was largely decreased compared to neurons preincubated with conditioned medium from neurons expressing α-syn^WT^-GFP (Supp Fig. 1 B). We also asked whether the conditioned medium from α-syn^WT^-expressing neurons could affect the frequency of CTB+ carriers when applied on the soma chamber of microfluidic device. However, no effect was observed (Figure 1 D, E and F), suggesting that the conditioned medium acted primarily on presynapses.

**Figure 1:**
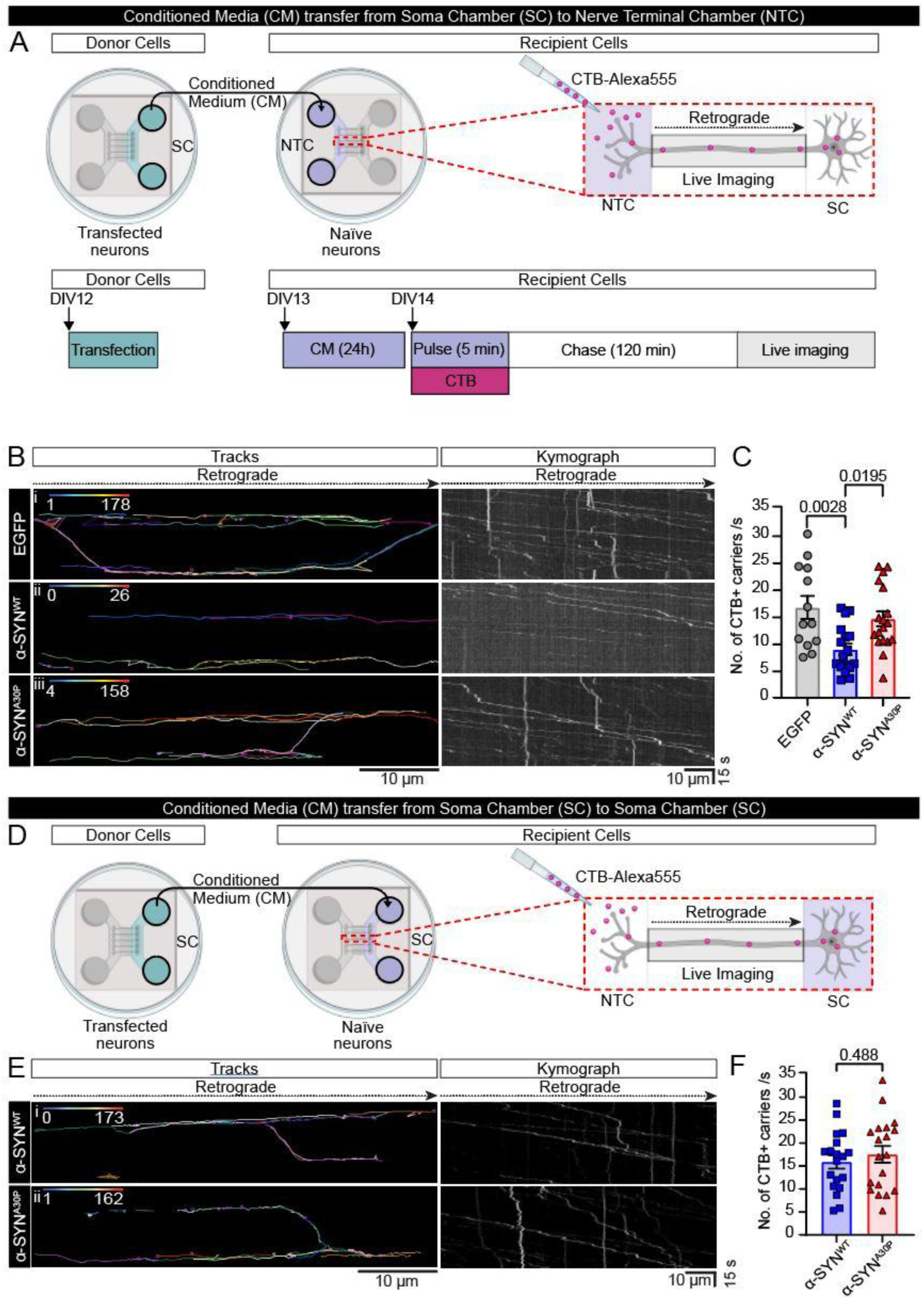
Conditioned medium from hippocampal neurons expressing α-syn^WT^-GFP downregulates axonal retrograde transport of CTB-positive carriers in recipient neurons. **A.** Illustration of the experimental workflow. Microfluidic chambers were seeded with hippocampal neurons extracted from brains of E16 mice embryos which were transfected with either EGFP, α-syn^WT^ -GFP or α-syn^A30P^ -GFP (donor device; DIV 12). After 24h the conditioned medium from the soma chamber (SC) of the donor device was added to the nerve terminal chamber (NTC) of the recipient device (70:30 ratio) containing untransfected naïve neurons. Following a 24h incubation, a 5-minute pulse of Cholera toxin subunit B conjugated to Alexa Fluor™ 555 (CTB-Af555; 50 ng/mL) in high potassium buffer was conducted followed by 3 washes of the CTB with low potassium buffer and a 2h chase with the conditioned medium before imaging of CTB-positive (CTB+) carriers undergoing retrograde transport in axons of the recipient device. Timelapse confocal imaging was performed on the discovery spinning disk at a 3Hz imaging rate. **B.** Representative track images and kymographs of CTB+ carriers undergoing retrograde axonal trafficking. Recipient neurons that received i. EGFP, ii. α-syn^WT^ -GFP, or iii. α-syn^A30P^-GFP. Left panels, Trackmate tracing of CTB tracks in representative movies (colour-coded for the number of CTB+ carriers), scale bar: 10 µm. Right panels, corresponding kymographs. Scale bar: x-axis, 10µm; y-axis, 15s. **C.** Quantification of the frequency of CTB+ carriers. Data from 2 independent experiments, n = 13-18 channels imaged. Results present the mean ± SEM, one-way ANOVA, Tukey’s test. **D.** Illustration of the experimental workflow. Microfluidic chambers were seeded with hippocampal neurons extracted from brains of E16 mice embryos which were transfected with either EGFP, α-syn^WT^ -GFP or α-syn^A30P^ -GFP (DIV 12). After 24h, the conditioned medium from the soma chamber (SC) of the donor device was added to the soma chamber (SC) of the recipient device (70:30 ratio) containing untransfected naïve neurons. Following a 24h incubation, a 5-minute pulse of Cholera toxin subunit B conjugated to Alexa Fluor™ 555 (CTB-Af555; 50 ng/mL) in high potassium buffer was conducted, followed by 3 washes of the CTB with low potassium buffer and a 2h chase with the conditioned medium before imaging of CTB-positive (CTB+) carriers undergoing retrograde transport in axons of the recipient device. Timelapse confocal imaging was performed on the discovery spinning disk at a 3Hz imaging rate. **E.** Representative track images and kymographs of CTB+ carriers undergoing retrograde axonal trafficking. Recipient neurons that received i. α-syn^WT^ -GFP, or ii. α-syn^A30P^-GFP. Left panels, Trackmate tracing of CTB tracks in representative movies (colour-coded for the number of CTB+ carriers), scale bar: 10µm. Right panels, corresponding kymographs. Scale bar: x-axis, 10µm; y-axis, 15s. **F.** Quantification of the frequency of CTB+ carriers. Data from 2 independent experiments, n= 19 channels imaged. Results present the mean ± SEM, unpaired Student t-test.

Since exosomal transmitted α-syn has been implicated in neuronal death (Emmanouilidou et al. 2010; Danzer et al. 2012; Ngolab et al. 2017), we reasoned that the effect of α-syn^WT^ conditioned medium could be mediated by the release of α-syn located in exosomes following the fusion of MVBs with the plasma membrane. To visualise exosomal release, we used a common exosomal marker, CD63, coupled to a pH sensitive green fluorescent reporter, pHluorin, which fluoresces only in a neutral environment, namely when exosomes are released from the acidic environment of the intraluminal MVB into the extracellular space (Miesenböck, De Angelis, and Rothman 1998; Piper and Katzmann 2007; Verweij et al. 2018). We transfected neurosecretory pheochromocytoma (PC12) cells with CD63-pHluorin and used Total Internal Reflection Fluorescence (TIRF) microscopy to image single exocytic events as previously performed (Verweij et al. 2018). We transfected neurosecretory pheochromocytoma (PC12) cells with CD63-pHluorin and used Total Internal Reflection Fluorescence (TIRF) microscopy to image single CD63-positive exocytic events as previously described (Verweij et al. 2018). Live imaging of exosomal release was visualised as discrete bursts in fluorescence intensity detected at the plasma membrane (Figure 2 A-E). Secretagogue stimulation using barium chloride promoted an increase in the number of exosomal release events (Figure 2 C). The duration of the fusion events was widespread, from less than a second to more than 500 seconds with an average signal duration of 34.12 seconds (Figure 2 D). Recently, the Heat Shock Protein 90 (Hsp90), a chaperone protein, was shown to control the fusion of MVBs with the neuronal plasma membrane, leading to the release of exosomes (Lauwers et al. 2018). Hsp90^MD7^ bearing 3 substitutions in its amphipathic helix, was uncovered in a screen for *Drosophila* proteins able to deform membranes and was found to prevent the fusion of MVBs with the plasma membrane (Figure 2 F). Co-transfection of PC12 cells with CD63-pHluorin and Hsp90^MD7^-mCherry significantly decreased the number of fusion events observed upon stimulation (68.25% decrease) compared to cells co-transfected with Hsp90^WT^ (Figure 2 F), thereby confirming the efficiency of Hsp90^MD7^ in blocking exosomal release from MVBs (Lauwers et al. 2018). Further, the duration of the detected exocytic events was not sensitive to Hsp90^MD7^ mutations (average of 35.57 seconds and 49.36 seconds for Hsp90^WT^ and Hsp90^MD7^ co-transfected cells respectively), suggesting that the few exocytic events detected were full and complete (Figure 2 I).

**Figure 2:**
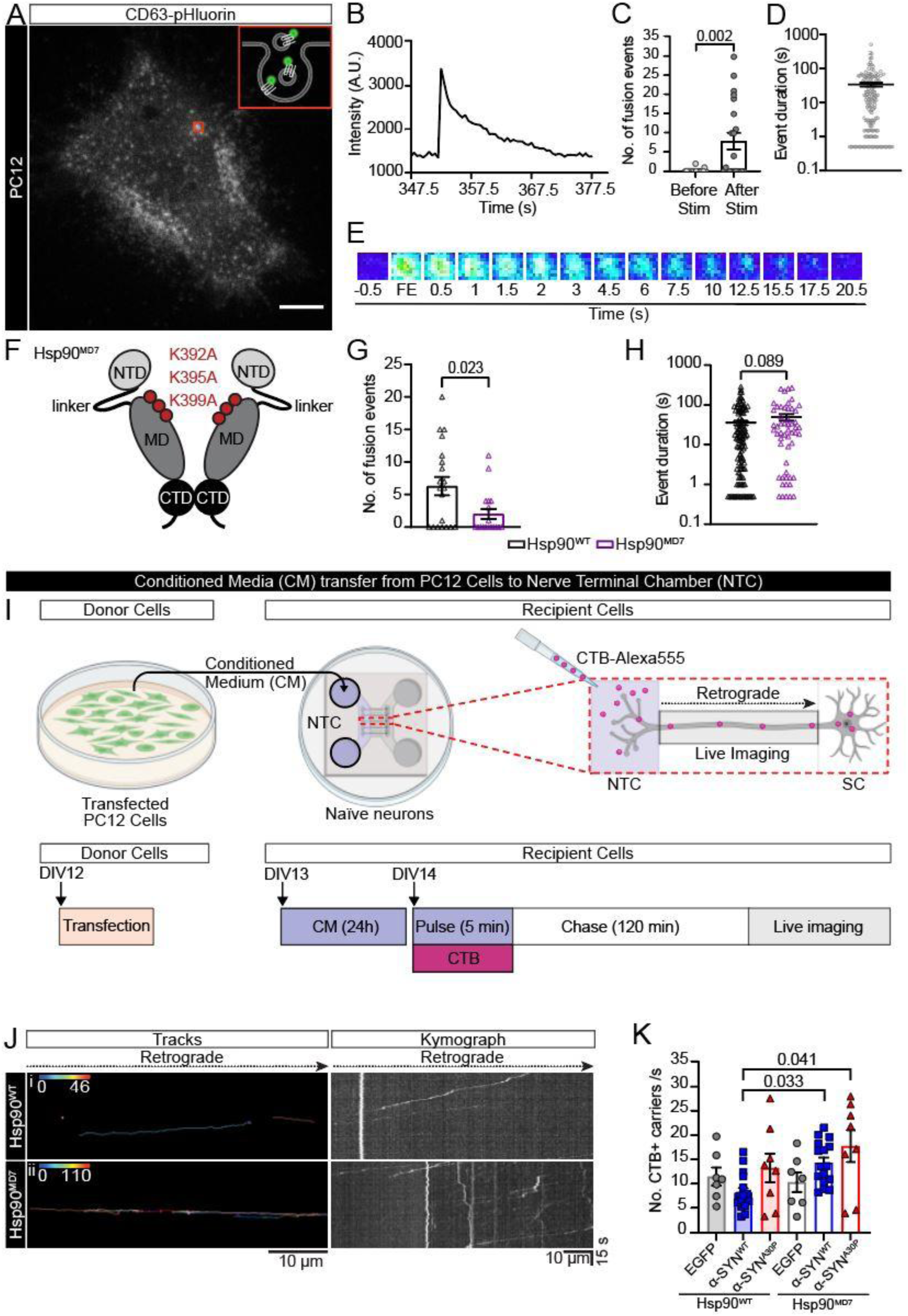
The downregulating effect of the conditioned medium is Hsp90-dependent. PC12 cells were co-transfected with an exosomal marker (CD63-pHluorin) and with either mCherry, Hsp90^WT^-mCherry or mutant Hsp90^MD7^-mCherry. 48h post transfection, TIRF imaging of exosomal release was performed at 2Hz on a Roper-iLas 2 microscope. A 10-minute timelapse was recorded with stimulation after 5 minutes using 2 mM barium chloride. **A.** A representative image of a PC12 cell membrane with CD63-pHluorin exocytic event outlined by a red box. **B.** Change in fluorescence intensity over time of the representative exocytic event, time in seconds. **C.** Number of CD63-pHluorin fusion events detected before and after stimulation with barium chloride in cells co-transfected with CD63-pHluorin and mCherry. Data from 4 independent experiments, n=21 cells. Results present the mean ± SEM. Wilcoxon paired test. **D.** Duration of CD63-pHluorin exocytic events (seconds) detected in cells co-transfected with mCherry. Data from 4 independent experiments, n=21 cells. **E.** Timelapse cartoon of the representative exocytic event, FE corresponding to the start of the fusion event and time in seconds. **F.** A representative image of a Hsp90^MD7^ dimer and the mutations done in the amphipathic helix. **G.** Number of CD63-pHluorin exocytic events detected in cells co-transfected with either Hsp90^WT^ or Hsp90^MD7^. Data from 4 independent experiments, n=17-19 cells. Results present the mean ± SEM. Mann-Whitney test. **H.** Duration of CD63-pHluorin exocytic events detected in cells co-transfected with either Hsp90^WT^ or Hsp90^MD7^, time in seconds. Data from 4 independent experiments, n=17-19 cells. Results present the mean ± SEM, Mann-Whitney test. **I.** Illustration of the experimental workflow. Microfluidic chambers were seeded with hippocampal neurons extracted from brains of E16 mice embryos. PC12 cells were co-transfected with either Hsp90^WT^-mCherry or Hsp90^MD7^-mCherry in combination with either EGFP, α-syn^WT^ -GFP or α-syn^A30P^-GFP. After 24h, the conditioned medium from PC12 cells was added to the nerve terminal chamber (NTC) of microfluidic devices (70:30 ratio) containing untransfected naïve neurons (DIV 13). Following a 24h incubation, a 5-minute pulse of Cholera toxin subunit B conjugated to Alexa Fluor™ 555 (CTB-Af555; 50 ng/mL) in high potassium buffer was conducted, followed by 3 washes of the CTB with low potassium buffer and a 2h chase with the conditioned medium before imaging of CTB-positive carriers (CTB+) undergoing retrograde transport in axons of the recipient device. Timelapse confocal imaging was performed on the discovery spinning disk at a 3Hz imaging rate. **J.** Representative track images and kymograph of CTB+ carriers undergoing retrograde axonal trafficking. Recipient neurons that received i. α-syn^WT^ -GFP with Hsp90^WT^-mCherry or ii. α-syn^WT^-GFP with Hsp90^MD7^-mCherry. Left panels, Trackmate tracing of CTB tracks in representative movies (colour-coded for the number of CTB+ carriers), scale bar: 10µm. Right panels, corresponding kymographs. Scale bar: x-axis, 10µm; y-axis, 15s. **K.** Quantification of the number of CTB+ carriers. 2 independent experiments with 7-18 channels imaged. Results present the mean ± SEM, Kruskal-Wallis test.

Having demonstrated that Hsp90^MD7^ mutations can decrease exosomal release from PC12 cells, we reasoned that co-expression of Hsp90^MD7^-mCherry with α-syn^WT^-GFP in these cells could reduce α-syn exosomal release and impact retrograde trafficking in recipient neurons incubated with the PC12 cells conditioned medium (70/30 vol/vol ratio). To test this, we harvested conditioned media of PC12 cells co-transfected with either Hsp90^WT^-mCherry or Hsp90^MD7^-mCherry, in combination with either EGFP, α-syn^WT^-GFP or α-syn^A30P^-GFP and applied them on recipient neurons to assess their effect on the retrograde transport of CTB (Figure 2 I). As anticipated, the decrease in the number of CTB+ carriers retrogradely transported in axons incubated with conditioned medium generated in α-syn^WT^-GFP/Hsp90^WT^-mCherry co-expressing PC12 cells, was lost and returned to control levels (EGFP/Hsp90^WT^) in axons incubated with α-syn^WT^/Hsp90^MD7^ conditioned medium (Figure 2 J-K). This result demonstrated that the downregulating effect of α-syn^WT^ conditioned medium on retrograde trafficking required Hsp90-dependent exosomal release from donor cells. Interestingly, α-syn^A30P^/Hsp90^MD7^ conditioned medium did not impact axonal transport (Figure 2 K). This suggests that the lack of downregulating effect of α-syn^A30P^ conditioned medium on retrograde transport does not stem from an altered exosomal release. The downregulating effect of conditioned medium from α-syn^WT^-GFP expressing neurons is therefore likely to be mediated by exosomal α-syn^WT^.

As α-syn is known to be secreted either as free membrane-less monomer/aggregate (El-Agnaf et al. 2003; Lee, Patel, and Lee 2005; Xie et al. 2022) or within different vesicles (Emmanouilidou et al. 2010; Ejlerskov et al. 2013; Minakaki et al. 2018; Yamada and Iwatsubo 2018), we reasoned that removing the free forms of α-syn^WT^-GFP should not impact on the ability of the conditioned medium to downregulate axonal retrograde transport. To test this hypothesis, we immunodepleted α-syn^WT^-GFP from the conditioned medium of PC12 cells co-transfected with α-syn^WT^-GFP and Hsp90^WT^-mCherry using anti-GFP trap beads (Supp Fig 2 A). We first checked that the conditioned medium was depleted from α-syn^WT^-GFP by Western Blot. As expected, α-syn-GFP was detected in the original conditioned medium and in the immunoprecipitated beads fraction (Supp Fig 3 B). Interestingly, we could not detect α-syn-GFP in the flowthrough suggesting that the secreted form consists primarily of free α-syn monomers/aggregates. The immunodepleted and original conditioned media were consequently applied onto the nerve terminal chamber of microfluidic devices (70/30 volv/vol ratio) seeded with naïve hippocampal neurons allowing separation of the cell body from nerve terminals to study axonal transport. After a 24-hour incubation with the conditioned media mix and a 5-minute pulse with CTB-Af555 diluted in HK^+^ buffer, devices were washed three times with LK^+^. Neurons were incubated back in the conditioned media mix for 2 hours before performing timelapse imaging of CTB-positive (CTB+) carriers undergoing retrograde transport in axons of recipient neurons. As anticipated, the number of CTB+ carriers retrogradely transported in recipient neurons incubated with the α-syn^WT^-GFP immunodepleted conditioned medium from α-syn^WT^-GFP/ Hsp90^WT^-mCherry co-expressing cells did not statistically change from the number of CTB carriers observed in neurons incubated with the original conditioned medium and was still significantly lower compared to the number of carriers in neurons incubated with conditioned medium from PC12 cells co-expressing α-syn^WT^-GFP/ Hsp90^MD7^-mCherry (Supp Fig 3 C). These results suggest that the exosomal form of α-syn is relatively minor but plays a key role in downregulating retrograde trafficking in recipient neurons.

**Figure 3.**
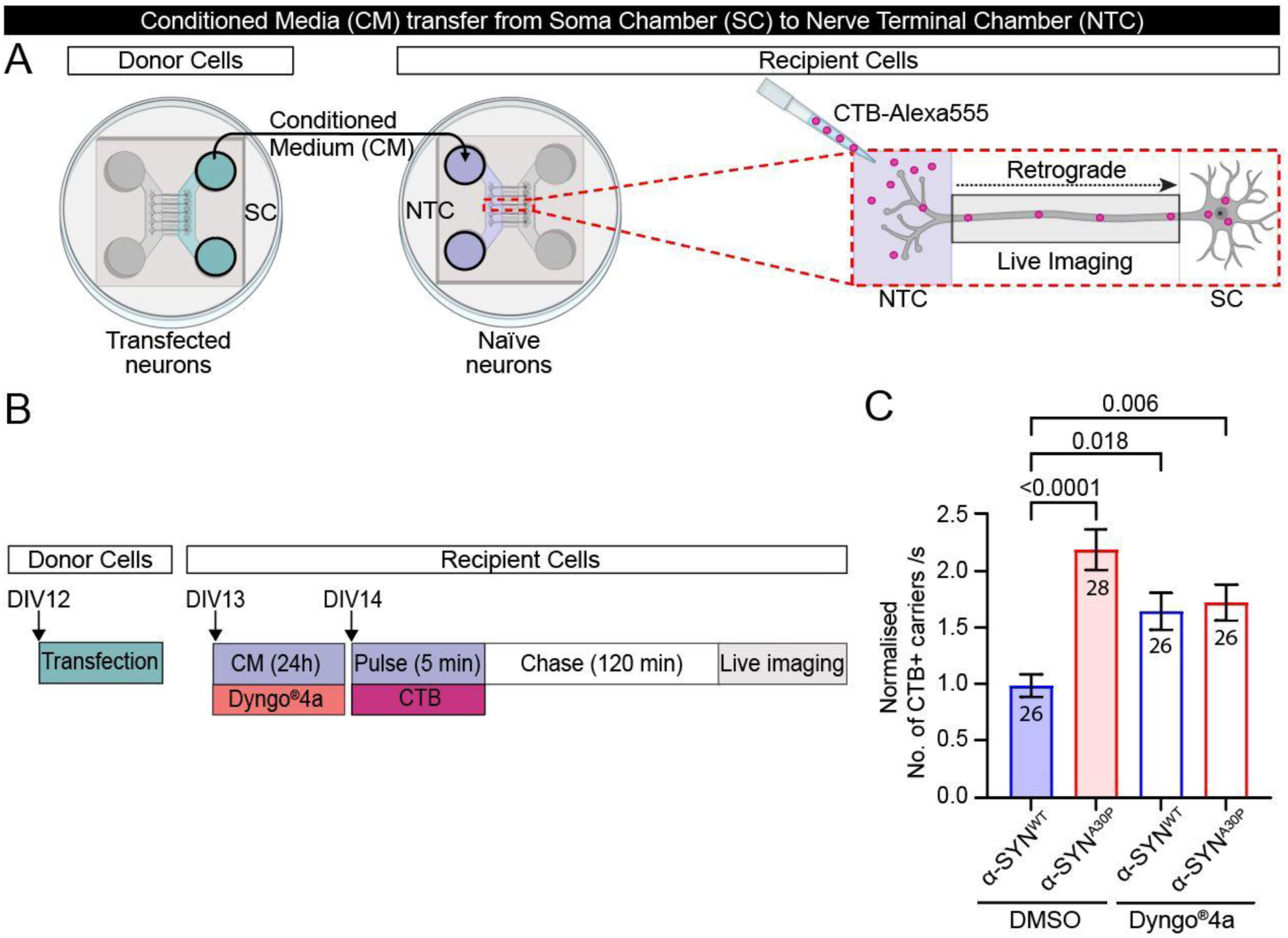
The downregulating effect of the conditioned medium in recipient neurons is dependent on endocytosis. A-B. Schematic of the experimental paradigm: Microfluidic chambers were seeded with hippocampal neurons extracted from brains of E16 mice embryos and neurons from the soma chamber (SC) were transfected with either α-syn^WT^-GFP or α-syn^A30P^-GFP (donor device; DIV 12). After 24h, the conditioned medium from the SC of the donor device was added to the nerve terminal chamber (NTC) (70:30 ratio) of the recipient device containing untransfected naïve neurons, in the presence of Dyngo^®^4a (30µM) or DMSO. The following day, a 5-minute pulse with Cholera toxin subunit B conjugated to Alexa Fluor™ 555 (CTB-Af555; 50 ng/mL) diluted in high potassium (HK^+^) buffer was conducted followed by 3 washes with low potassium (LK^+^) buffer to remove the excess of CTB. Devices were then incubated for 2h at 37°C with Dyngo^®^4a free-conditioned medium before imaging of CTB-positive carriers undergoing retrograde transport in axons of the recipient devices. Timelapse confocal imaging was performed on the Discovery spinning disk at a 3Hz imaging rate. **C.** Quantification of the number of CTB+ carriers. Data from 3 independent experiments, n=26-28 channels, one-way ANOVA followed by Tukey’s multiple comparison test, mean ± SEM.

To verified that α-syn^WT^-GFP was indeed released following MVB fusion events, we carried out further experiments in hippocampal neurons using CD63-pHuji (red fluorescent version of pHluorin) as an exosomal fluorescent marker. The goal was to visualise α-syn-GFP release from MVBs by assessing the change in fluorescence in the green and red channels in neurons transfected with CD63-pHuji and either EGFP, α-syn^WT^-GFP, or PD-linked mutant α-syn^A30P^-GFP (Supp Fig. 3 A-C). We detected CD63-pHuji exocytic events in the red channel on the cell body of hippocampal neurons, that were typically characterised by a rapid drop in fluorescent intensity (Supp Fig 3 B-D). This was associated in 40% of the cases with a slow increase in α-syn^WT^-GFP observed in the green channel (Supp Fig 3 B, C and E) reflecting the slow release of α-syn^WT^-GFP exosomes. The number of fusion events was increased upon depolarisation with high K^+^ buffer (Supp Fig 3 F) paralleling the effect detected in neurosecretory PC12 cells (Figure 2). Interestingly, there was no statistical difference between α-syn^WT^ and α-syn^A30P^ release events in terms of numbers, percentage of events containing α-syn-GFP and changes in fluorescence intensity, suggesting that both WT and mutant forms of α-syn were equally released following MVB fusion (Supp Fig 3 F-H). These results demonstrate that α-syn is released from MVB and that both the WT and mutant forms are equally released. This raises the question of the origin of the negative regulation of vesicular trafficking afforded by the conditioned medium from α-syn^WT^-GFP expressing neurons.

Considering that the conditioned medium from α-syn^A30P^-GFP expressing neurons had no impact on retrograde trafficking, and that its exosomal release from donor neurons was equivalent to that of α-syn^WT^-GFP (Supp Fig 3), we reasoned that these effects could stem from the differential uptake of α-syn^WT^-GFP or α-syn^A30P^-GFP in recipient neurons. The uptake in neuroblastoma cells of exosomes isolated from patients with Lewy Body Dementia, is energy dependent and mediated by dynamin-dependent endocytosis (Ngolab et al. 2017). Furthermore, the integrity of exosomes is essential for the uptake of exosomal α-syn (Alvarez-Erviti et al. 2011; Danzer et al. 2012). We therefore used a selective dynamin inhibitor (Dyngo^®^4a), known to block the major types of endocytosis in neurons (Newton et al. 2006 PNAS; Gormal et al. 2017, Nguyen et al. 2013; Harper et al. 2011 Harper et al. 2013), to test whether the negative regulation of axonal retrograde transport afforded by the conditioned medium was sensitive to dynamin inhibition in recipient neurons. We transfected hippocampal donor neurons with α-syn^WT^-GFP or α-syn^A30P^-GFP, harvested the conditioned media after 24 hours and incubated recipient neurons from the nerve terminal chamber, in the presence of Dyngo4a® (Figure 3 A-B) for another 24 hours. Following a brief wash with fresh conditioned medium to remove the dynamin inhibitor, we pulsed the nerve terminal chamber with CTB-Af555 diluted in HK^+^ buffer for 5 minutes, washed chambers three times with LK^+^ buffer and put back the conditioned medium for a 2-hour chase at 37°C. Timelapse imaging and quantification of CTB-AF555 retrograde transport in axons of recipient neurons was then performed (Figure 3 A-C). The number of CTB+ carriers was increased in recipient neurons exposed to α-syn^WT^-GFP conditioned medium and Dyngo^®^4a compared to neurons exposed to α-syn^WT^-GFP conditioned medium only, demonstrating the downregulation of retrograde trafficking relies on dynamin-dependent endocytosis (Figure 3 C). The conditioned medium from the PD-linked mutant α-syn^A30P^-GFP was used as a negative control, as it did not downregulate retrograde trafficking (Figure 3 C and Figure 1 B-C). Our findings demonstrate that dynamin-dependent endocytosis is required in nerve terminals for α-syn^WT^-GFP conditioned medium to downregulate retrograde trafficking in recipient neurons.

“Seeing is believing”. The effect of exosomal α-syn contained in the conditioned medium requires dynamin-dependent uptake in recipient nerve terminals. To confirm that exosomal α-syn is indeed uptaken by recipient neurons, we asked whether we could image it in recipient neurons. However, released exosomal α-syn is a relatively minor fraction of extracellular α-syn (Supp Fig 2 B). Further, the size of exosomes typically ranges from 30 to 100 nm, which is below the diffraction limit of light (Théry, Zitvogel, and Amigorena, 2002). For these reasons, we turned to single molecule super-resolution microscopy to be able to detect sparce molecules of transmitted α-syn in recipient neurons. Recipient hippocampal neurons (DIV12-14) were incubated for 24 hours with conditioned medium from neurons expressing either mEos2 alone, α-syn^WT^-mEos2, or α-syn^A30P^-mEos2 (Figure 4 A). Neurons were pulsed for 5 minutes with CTB-Af647 in HK^+^ buffer, washed 3 times in LK^+^ buffer before a 2-hour incubation back with the conditioned medium. We then performed single-particle tracking PhotoActivated Localization Microscopy (sptPALM) of transmitted mEos2, α-syn^WT^-mEos2 or α-syn^A30P^-mEos2, which allows live tracking of individual molecules in living cells (Manley et al. 2008). As expected, mEos2, α-syn^WT^-mEos2 and α-syn^A30P^-mEos2 were detected in the axons of recipient neurons that were positive for CTB-Af647 (Figure 4 B). Data analysis following tracking of individual trajectories was carried out and their nanoscale mobility was derived from the mean square displacement (MSD). The number of trajectories, their duration and the mobile/immobile ratio were also determined. Interestingly, the mobility of α-syn^WT^-mEos2 was significantly reduced compared to that of mEos2 and α-syn^A30P^-mEos2 (Figure 4 B) and displayed prominent periodicity (Figure 4 C). α-syn^WT^-mEos2 molecules displayed a clustered behaviour with a cluster size of 177 ± 5 nm, distance between clusters of 391 ± 23 nm and a periodicity of 393 ± 26 nm. Notably, recipient neurons contained less transmitted α-syn^WT^-mEos2 compared to mEos2 and α-syn PD-linked mutant suggesting a potential effect of transmitted α-syn^WT^-mEos2 on the uptake mechanism. However, the percentage of uptake was not significant between conditions (Figure 4 D). No significant difference was seen in trajectory duration or in mobile/immobile ratio Furthermore, no significant difference was seen in trajectory duration nor in the mobile/immobile ratio (Figure 4 D).

**Figure 4:**
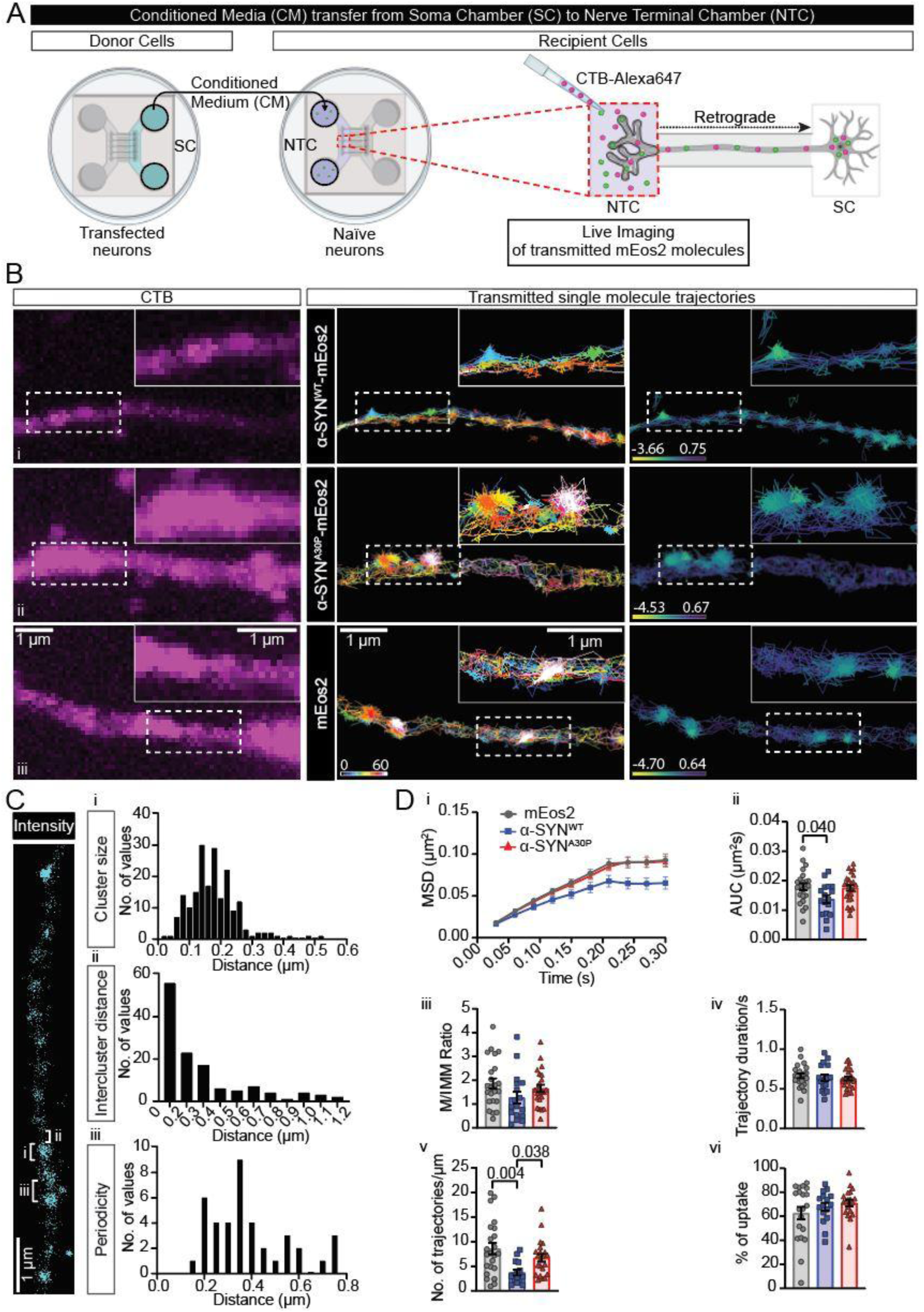
Single molecule imaging of transmitted α-syn in recipient neurons reveals distinct mobilities and a periodicity. **A.** Detailed schematic of the experimental design: Microfluidic chambers were seeded with hippocampal neurons extracted from brains of E16 mice embryos and neurons from the soma chamber (SC) of the devices were transfected with either mEos2, α-syn^WT^-mEos2 or α-syn^A30P^-mEos2 (donor device; DIV 11-13). After 24h, the conditioned medium from the SC of the donor device was added to the nerve terminal chamber (NTC) (70:30 ratio) of the recipient device containing untransfected naïve neurons. The following day, a 5-minute pulse with Cholera toxin subunit B conjugated to Alexa Fluor™ 647 (CTB-Af647; 50 ng/mL) diluted in high potassium (HK^+^) buffer was conducted followed by three washes with low potassium (LK^+^) buffer to remove the excess of CTB. Devices were then incubated back with the conditioned medium for 2h at 37°C before imaging of CTB-positive carriers undergoing retrograde transport in axons and imaging of mEos2 molecules in axons of the recipient devices. Timelapse imaging was performed on the Elyra microscope. **B.** Left panel, images of incorporated CTB-Af647 in recipient neurons incubated with conditioned medium from neurons expressing allowing to draw region of interest around axons i. α-syn^WT^-mEos2, ii. α-syn^A30P^-mEos2 or iii. mEos2. Middle panel, sptPALM super-resolved trajectory images of either, i. α-syn^WT^-mEos2, ii. α-syn^A30P^-mEos2 or iii. mEos2 in recipient neurons; the colour gradient of the calibration bar indicates the time of detection of trajectories (seconds): dark blue corresponding to a molecule detected at the beginning of movie acquisition and white corresponding to a molecule detected at the end of the acquisition. Right panel, super-resolved NASTIC diffusion coefficient images of either, i. α-syn^WT^-mEos2, ii. α-syn^A30P^-mEos2 or iii. mEos2 in recipient neurons where each trajectory is coloured according to the value of the diffusion coefficient (log10[μm^2^s^−1^]) for the first four time points, yellow corresponding to more immobile molecules whilst dark blue to mobile ones. Scale bar = 1 µm. **A.** Periodicity analysis of α-syn^WT^-mEos2 molecules in recipient neurons; analysis was carried out on the intensity map as displayed in a representative image on the left panel, scale bar = 1 µm, with analysis of i. α-syn^WT^-mEos2 cluster size, ii. Distance between clusters and iii. Periodicity of clusters. **B.** Mobility analysis of mEos2 molecules for all 3 conditions in recipient neurons (mEos2, α-syn^WT^-mEos2 or α-syn^A30P^-mEos2), i. Graph of the mean square displacement (MSD, µm^2^), ii. Graph of the area under the MSD curve (AUC, µm^2^s), one-way ANOVA followed by Tukey’s multiple comparison test, mean ± SEM. iii. Graph of the mobile/immobile ratio, Kruskal-Wallis test followed by Dunn’s multiple comparison test, mean ± SEM iv. Graph of the duration of trajectories (seconds), one-way ANOVA followed by Tukey’s multiple comparison test, mean ± SEM. v. Graph of the number of trajectories per µm, Kruskal-Wallis test followed by Dunn’s multiple comparison test, mean ± SEM. Data from 10 independent experiments, n= 17-24 axons.

## Discussion

In this study, we have established that conditioned medium from donor hippocampal neurons expressing wild type α-syn (α-syn^WT^), but not PD-mutant A30P (α-syn^A30P^) or EGFP was found to downregulate retrograde trafficking of CTB in recipient neurons. This effect relied on an Hsp90-dependent release of exosomes. We further characterised the release of exosomal α-syn from hippocampal neurons and its generic component. In addition, we demonstrated that the uptake of α-syn containing exosomes was also generic, relying on a dynamin-dependent internalisation. Super-resolution single molecule imaging of transmitted α-syn allowed us to confirm the generic nature of the transmission to recipient neurons but also highlighted a surprising periodic nanoscale organisation of transmitted α-syn^WT^ and its lower mobility suggestive of an action in recipient nerve terminals and axons.

Parkinson’s disease (PD) is characterised by an initial loss of dopaminergic neurons and there is increasing evidence of synaptic dysregulation preceding neurodegeneration (Kramer and Schulz-Schaeffer 2007; Chung et al. 2009; Scott et al. 2010; Tagliaferro and Burke 2016; Soukup, Vanhauwaert, and Verstreken 2018). Soukup, Vanhauwaert, and Verstreken 2018). Indeed, α-syn aggregates have been found to accumulate in the presynaptic compartment with noticeable axon terminals degeneration in the brain of PD and DLB patients (Galvin et al. 1999; Kramer and Schulz-Schaeffer 2007). Also, addition of exogenous α-syn fibrils to neurons leads to the formation of inclusions, firstly in axons before propagation to the cell body (Volpicelli-Daley et al. 2011). Long-range retrograde transport is crucial for neuron survival (Heerssen, Pazyra, and Segal 2004; Wang et al. 2016) and seems to be impacted in synucleinopathies. Indeed, in sporadic cases of PD and in a rat model of synucleinopathy, levels of axonal transport proteins have been shown to be decreased (Chung et al. 2009; Chu et al. 2012). Here, we show that conditioned medium from α-syn^WT^ expressing neurons specifically downregulates retrograde trafficking of CTB in recipient neurons. This effect on retrograde transport observed with conditioned medium from α-syn^WT^ expressing neurons, which is lost with the PD mutant A30P, could potentially function to regulate the frequency of signalling endosomes that are transported from the presynapse to the cell body and thus affect neuronal survival. Our finding that neurons exposed to α-syn^WT^ conditioned medium presented an increased colocalization of CTB with TrkB compared to PD mutant α-syn^A30P^ corroborates previous findings that exogenous α-syn (fibrils in the study) can alter the transport of TrkB in hippocampal neurons (Volpicelli-Daley et al. 2014). It can be hypothesised that the deregulation of the transport of signalling endosomes in recipient neurons exposed to exogenous α-syn could be one of the mechanisms involved in neuronal death. The fact that conditioned medium from α-syn^WT^ expressing neurons impacted CTB retrograde transport in recipient neurons only when it was added to the nerve terminal chamber of microfluidic devices, but not to the soma chamber, is consistent with the hypothesis of neuronal death stemming from an early dysfunction at the presynapse. It suggests α-syn can enter nerve terminals and exert its effect there. This is in line with *in vivo* experiments carried out in mice and in non-human primates, where propagation of intracerebrally injected α-syn was found to be transsynaptic (Luk et al. 2012; Recasens et al. 2014; Okuzumi et al. 2018).

α-syn is a cytoplasmic protein and is known to be secreted from neurons either in a free form (Lee, Patel, and Lee 2005; Emmanouilidou et al. 2010; Jang et al. 2010, Ejlerskov et al. 2013; Yamada and Iwatsubo 2018; Xie et al. 2022) or within extracellular vesicles (Lee, Patel, and Lee 2005; Emmanouilidou et al. 2010; Jang et al. 2010; Danzer et al. 2012; Ejlerskov et al. 2013; Minakaki et al. 2018). Amongst the vesicular forms of secreted α-syn, exosomal α-syn represents a minority of the α-syn contained in conditioned medium (Alvarez-Erviti et al. 2011; Hasegawa et al. 2011; Delenclos et al. 2017), but it is nonetheless toxic for surrounding cells, with recipient neuroblastoma cells being less viable (Emmanouilidou et al. 2010; Danzer et al. 2012). This difference in α-syn secretion was also found in our hands in PC12 cells, where the fraction containing free α-syn gave a strong signal as opposed to no signal on the Western Blot for the fraction containing the vesicular forms of α-syn. The absence of signal for the vesicular fraction could be due to a low amount of vesicular α-syn as we used small volumes of conditioned medium (1 mL) without prior concentration step. Harvesting a bigger volume of media, followed by a concentration step might be required to obtain a detectable signal. Characterisation of exosomal release in PC12 cells using CD63-pHluorin gave similar results to what was documented for other cell lines. Exosomal release was reflected by a sharp increase in green fluorescence, in agreement with previous literature (Verweij et al. 2018; Bebelman et al. 2020; Mahmood et al. 2023). The number of fusion events was variable between cells, which was reported as well (Verweij et al. 2018; Bebelman et al. 2020). The number of fusion events per minute observed in cells expressing mCherry was similar to that observed in HUVEC cells (Bebelman et al. 2020). Majority of fusion events observed in PC12 cells were indeed multivesicular body (MVB) fusions with the plasma membrane as the signal duration was longer than 2 seconds (Verweij et al. 2018; Bebelman et al. 2020). However, the average duration in mCherry co-transfected cells was slightly shorter compared to what was described in Hela cells (34.12 seconds in PC12 cells versus 106.55 seconds in Hela cells). It could be that MVBs in PC12 cells have different size distribution and composition compared to HeLa cells leading to different fusion kinetics. It is worth noting that analysis of exosomal release was here done manually. To then assess the effect of Hsp90 mutations in PC12 cells, cells were co-transfected with CD63-pHluorin and Hsp90^MD7^-mCherry. Analysis revealed a 68% decrease in exosomal release compared to Hsp90^WT^-mCherry co-transfected cells. The effect of Hsp90 mutations on exosomal release was slightly stronger in PC12 cells compared to what was observed in the neuromuscular junction of *Drosophila* (50% decrease) (Lauwers et al. 2018). This could be explained by the overexpression of Hsp90^MD7^ in PC12 cells as opposed to expression of Hsp90^MD7^ at endogenous levels in *Drosophila*. Interestingly, when conditioned medium from α-syn^WT^-GFP/ Hsp90^MD7^-mCherry co-transfected cells was applied to recipient neurons, the downregulating effect on retrograde transport was abolished, confirming the implication of exosomes in the effect. The fact that the fraction containing only free α-syn^WT^ did not decrease the number of CTB+ carriers in recipient neurons strengthens this hypothesis. However, to confirm that exosomes containing α-syn^WT^ are specifically responsible of this effect, and not another type of extracellular vesicles containing α-syn^WT^, we would need to isolate exosomes from the conditioned medium and directly apply them on neurons to determine if the effect on the vesicular long-range retrograde trafficking observed with the conditioned medium is replicated with the exosomes containing α-syn^WT^.

Afterwards we investigated α-syn exosomal release in hippocampal neurons co-transfected with CD63-pHuji and α-syn-GFP, expecting to see a similar burst of fluorescence in the red channel that was seen with CD63-pHluorin in PC12 cells. Although, we instead observed a sharp decrease in red fluorescence. This could be potentially due to differences in the pH of the MVBs in neurons. Indeed, the average size of MVBs in neurons was bigger (diameter of 1.29 µm) compared to the size range reported in the literature where MVBs range from 0.25 to 1 µm (Von Bartheld and Altick 2011) with a majority around 0.4 to 0.6 µm (Altick et al. 2009; Ostrowski et al. 2010; Bebelman et al. 2020; Mahmood et al. 2023). Due to their increased size, MVBs might not have been subject to an acidification up to pH 5.5. This could have led to an incomplete quenching of the fluorescence (Shen et al. 2014) and explain the lack of initial burst of fluorescence upon fusion of MVBs with the plasma membrane. The signal duration for CD63-pHuji was shorter compared to CD63-pHluorin, but both experiments were done in different cells and parameters of exosomal release have been shown to be cell dependent (Bebelman et al. 2020). It is worth noticing that to the best of our knowledge, it is the first time exosomal release kinetics are described in PC12 cells and in hippocampal neurons using live imaging of CD63-pHluorin/CD63-pHuji. Interestingly, the increase in green fluorescence, corresponding to α-syn-GFP release, was always concomitant to a decrease in red fluorescence, suggesting that, α-syn-GFP was indeed released in large packets of exosomes. Although α-syn was described to also be released from other types of vesicles (Choi, Park, and Park 2021), it was not the case here as the increase in green fluorescence (α-syn-GFP release) was never observed on its own. Finally, we observed that α-syn exosomal release from neurons was the similar for WT, mutant A30P and was also occurring with EGFP. The generic process of exosomes release, with exosomes containing EGFP alone, was previously been described in the literature (Danzer et al. 2012; Brás et al. 2022).

Since the release of exosomal α-syn-GFP was similar for WT and PD-linked mutant, we went to see if the uptake might be different using a dynamin inhibitor (Dyngo^®^4a). α-syn fibrils (Lee et al. 2008; Hansen et al. 2011; Konno et al. 2012; Reyes et al. 2015; Masaracchia et al. 2018; Sengupta et al. 2020), and exosomal α-syn (Ngolab et al. 2017) have been shown to be uptaken in neurons via endocytosis. Our findings corroborate this, as incubating conditioned medium from neurons expressing α-syn^WT^ with a selective dynamin inhibitor (Dyngo^®^4a) prior to incubation with neurons and during the treatment with the conditioned medium, abolished the downregulation of CTB retrograde transport in recipient neurons observed with only the conditioned medium. One paper investigating internalisation of exosomal α-syn in neuroblastoma H4 cells found the uptake was not dependent on caveolin nor clathrin-mediated endocytosis or heparan sulphate proteoglycan receptor-mediated endocytosis (Delenclos et al. 2017). This result might appear contradicting with ours as clathrin-dependent endocytosis requires dynamin, however, several parameters in their study were different from our experimental design. They used Chlorpromazine which is a cationic amphiphilic drug thought to prevent the formation of clathrin-coated pits (Vercauteren et al. 2010) and they applied isolated exosomal α-syn instead of conditioned medium to neuroblastoma cells. The endocytosis of exosomal α-syn might involve different pathways in those cells compared to ones in neurons.

We then thought to image transmitted α-syn within hippocampal neurons using super resolution microscopy. We were able to detect exogenous α-syn-mEos2 in recipient neurons as well as mEos2 alone. The observation that α-syn and fluorophore proteins alone can transfer from one cell to another was previously described in neuroblastoma and neurosecretory cell lines (Hansen et al. 2011; Konno et al. 2012; Reyes et al, 2015). The co-culturing of SH-SY5Y neuroblastoma cells expressing α-syn-mCherry or mCherry alone with neurosecretory PC12 cells led to low transmission of proteins, with only a small fraction (4%) of neurosecretory PC12 cells containing α-syn inclusions and even less inclusions of mCherry (less than 0.5%) even after 72h of co-culturing (Konno et al. 2012). This low transmission was also observed in N2a neuroblastoma cells with 8% of double labelled cells after 5 days of culture (Reyes et al. 2015). In our case, we saw a large amount of α-syn-mEos2 and mEos2 alone moving within axons of recipient neurons after a 24-hour incubation with conditioned medium, as did another study using α-syn fibrils in cortical neurons seeded in microfluidic devices (Freundt et al. 2012). It might be that neurons have a higher ability of uptake compared to cell lines, or that the degradation process of the proteins is slower, leading to a higher amount within neurons.

From the imaging data, we were able to determine the mobility of transmitted α-syn within neurons and found that α-syn^WT^ had a lower mobility in axons compared to control and PD mutant α-syn^A30P^. To the best of our knowledge, it is the first time that live imaging of transmitted α-syn undergoing axonal transport in neurons has been done with the ability to distinguish single molecules of α-syn and study their confinement within neurons. Transport and mobility of endogenous α-syn has been studied using various methods showing that α-syn was transported via the slow components (Li et al. 2004; Saha et al. 2004; Roy et al. 2007; Roy et al. 2008; Yang et al. 2010). Several studies compared the mobility of α-syn^WT^ and mutants A30P and A53T. In two of them the velocity of axonal α-syn was decreased only for α-syn^A53T^ and not for α-syn^A30P^ (Li et al. 2004; Yang et al. 2010). A third study conducted in cortical neurons transfected with α-syn^WT^, α-syn^A30P^ and α-syn^A53T^ found that axonal transport was decreased for both mutants at early time points after transfection (3-4 hours) but then evened out at later time points to the level of α-syn^WT^ (Saha et al. 2004). Even if those studies seem to present contradictory results to ours, it is important to keep in mind that those studies looked at the velocity of α-syn which cannot be directly compared to the mean square displacement used in our experiment. Furthermore, those studies investigated α-syn expressed within neurons and not the transport of exogenous α-syn. One study looked at kinetics of axonal transmitted α-syn in cortical neurons exposed to α-syn fibrils (Freundt et al. 2012) and found they were consistent with slow-component-b transport and characterised by steady movement coupled to periods of pauses and oscillations observed previously for α-syn (Roy et al. 2007; Roy et al. 2008). mEos2 and α-syn-mEos2 displayed those behaviours in hippocampal neurons. The difference in α-syn^WT^ mobility we observed could be due to a specific interaction of α-syn^WT^ with cytoplasmic components, which potentially could be responsible of the downregulation of retrograde trafficking, only observed with conditioned medium from neurons expressing α-syn^WT^. The periodicity of α-syn^WT^ clustering in axons, which was not as obvious for the mutant A30P or mEos2 alone, supports this hypothesis. One of transmitted α-syn^WT^’s target could be endogenous α-syn since α-syn fibrils (Volpicelli-Daley, Luk, and Lee 2014; Fares et al. 2016; Karpowicz et al. 2017) and exosomal α-syn (Stuendl et al. 2016; Ngolab et al. 2017; Grey et al. 2015) have been shown to interact with endogenous α-syn in neurons. Because in our experiment we added conditioned medium of neurons expressing α-syn-mEos2, which contains free and exosomal α-syn, it is not possible to conclude that exosomal α-syn is directly responsible of downregulating retrograde transport, and that this effect is mediated by an interaction with endogenous α-syn. To prove this point, further experiments should be carried out by adding isolated exosomal α-syn to recipient neurons. The fact that we detected less trajectories for α-syn^WT^ compared to mutant A30P and mEos2 alone could be due to α-syn’s effect on endocytosis. Indeed, addition of monomeric α-syn^WT^ to slices containing rat calyces of Held presynaptic terminals and postsynaptic principal cells of the medial nucleus of the trapezoid body led to decreased endocytosis (Eguchi et al. 2017). Because we incubated neurons with conditioned medium for 24 hours, it might be that α-syn^WT^ from the conditioned medium reduced endocytosis in neurons leading to less uptake of α-syn^WT^ compared to the control and α-syn^A30P^. Further experiments need to be done to understand mechanisms involved in the effect of transmitted α-syn^WT^ and its long-term impact on neurons.

Here we unravelled, for the first time, the ability of transmitted exosomal α-syn^WT^ to downregulate retrograde trafficking in recipient neurons. We also demonstrated that this regulation is lost in a pathological context with PD mutant α-syn^A30P^. As retrograde trafficking is essential for neuronal survival, this loss of effect of exosomal α-syn^A30P^ could participate to α-syn pathology. Those results highlight the complexity of α-synucleinopathies where both exosomal and membrane-less transmitted α-syn play a crucial role in neurodegeneration.

## Materials and Methods

### Animal research ethics

All experiments using hippocampal neurons were performed on embryonic day 16 (E16) C57BL6/J mice brains according to the guidelines of the Australian Code for the Care and Use of Animals for Scientific Purposes and were approved by the Animal Ethics Committee from the University of Queensland (QBI/254/16/NHMRC and QBI/244/19/NHMRC).

### Hippocampal Neuronal Culture and transfection

Murine hippocampal neurons were dissected from E16 C57BL6/J mice as previously described (Joensuu et al. 2017). In brief, dissection of brains was performed in ice-cold dissection media (Milli-Q water, 1X Hank’s balanced salt solution (14065-056, Gibco), 10mM HEPES (15630080, Gibco), 100 units/mL-100 ug/mL Penicillin/Streptomycin (15140122, Gibco)). Trypsinisation of isolated hippocampi was done by adding 30 µL of Trypsin (15090046, Gibco) followed by 10 minutes incubation at 37°C. Digestion was stopped by adding 150 µL of heat-inactivated horse serum (HS) (26050088, Gibco) and 50 µL of 1% DNAse I (D5025-375KU, Sigma-Aldrich) followed by another 10-minute incubation at 37°C. After trituration of the sample and centrifugation for 7 minutes at 15000 rpm, neurons from the pellet were resuspended in Plating medium (5% foetal bovine serum (FBS) (10099141, Gibco), 100 units/mL-100 ug/mL Penicillin/Streptomycin (15140122, Gibco), 1X GlutaMax^TM^ (35050061, Gibco) in Neurobasal^TM^ medium (21103-049, Gibco) and plated on either glass-bottom dishes (D29-20-1.5-N, Cellvis) or on custom microfluidic devices (see section “microfluidic devices manufacturing”) at a density of 80,000-100,000 for glass bottom dishes, and respectively 5,000 and 50,000 for the nerve terminal and soma chambers of microfluidic devices (Joensuu et al. 2017). Prior to plating, dishes were coated with 1 mg/ml Poly-L-lysine (PLL) (P2636, Sigma-Aldrich) diluted in borate buffer (120 mM boric acid (B0394, Sigma-Aldrich) and 70 mM di-sodium tetraborate decahydrate (SO0707, Scharlau) and incubated overnight at 37°C before being washed three times with ultra-pure distilled water (10977-015, Invitrogen). After 3-4h, plating medium was replaced by Culture medium (1X B-27 Supplement (17504-044, Gibco), 1X GlutaMax^TM^ (35050061, Gibco) and 100 units/mL Penicillin with 100 ug/mL Streptomycin (15140122, Gibco) in phenol-free Neurobasal^TM^ medium (12348-017, Gibco)). Neurons were kept in the culture medium at 37°C/5% CO_2_ up to 20 days *in vitro* (DIV 20). 48h prior to imaging, hippocampal neurons were transfected with plasmids (list provided in the section “Plasmids and Fluorescent markers”) using Lipofectamine^TM^ 2000 (11668-019, Invitrogen) accordingly to the manufacturer’s protocol.

### PC12 Cell Culture and transfection

Rat pheochromocytoma cells (PC12) were cultured as previously described (Gormal et al. 2020). In brief, cells were grown in Culture medium (Dulbecco’s Modified Eagle Medium supplemented with 7% heat-inactivated HS (26050088, Gibco), 7% of heat-inactivated FBS (10099141, Gibco) and 0.85% Glutamax™ (35050061, Gibco)) at 37°C/5% CO_2_ in 75mm^2^ plastic culture flasks (156499, Thermofisher). 24h prior to transfection, cells were replated in 35mm Nunc EasyDish^™^ plastic dishes (150460, ThermoFisher) or in 6-well plates (92006, TPP or 3516, Cornstar). Transfection was carried out following the manufacturer’s protocol using the kit Lipofectamine LTX™/Reagent Plus™ (15338-100, Invitrogen) mixed with plasmids (list provided in the section “Plasmids and Fluorescent markers”) in OPTIMEM™ (31985070, Gibco). After 24h, either the conditioned medium was collected for immunodepletion and treatment of hippocampal neurons or cells were replated for imaging in glass-bottom dishes (D29-20-1.5-N, Cellvis) previously coated with 0.1 mg/ml Poly-D-lysine (P0899, Sigma-Aldrich) diluted in 1X phosphate-buffered saline solution (PBS) overnight and washed 3 times with ultrapure distilled water (10977-015, Invitrogen).

### Microfluidic device manufacture

Microfluidic devices were custom-made to fit inside a glass-bottom dish (D29-20-1.5-N, Cellvis) as previously described (Joensuu et al. 2017). Microfluidic devices allow separation of axons from cell bodies to facilitate the study of axonal transport (Taylor et al. 2005). Upon receipt, devices were rinsed one time with ultra-pure distilled water (10977-015, Invitrogen) followed by a coating step with PLL and seeding of hippocampal neurons in the devices (both steps are detailed in the section “Hippocampal Neuronal Culture and transfection” and in Joensuu et al. 2017).

### Neuronal treatment, stimulation, and nerve terminal labelling with CTB

On DIV 11-13, hippocampal neurons seeded in the soma chamber of microfluidic devices were transfected using Lipofectamine^TM^ 2000 (11668-019, Invitrogen) accordingly to the manufacturer’s protocol. Used plasmids are listed in the section “Plasmids and fluorescent markers”. After 24h, the conditioned medium from the soma chamber was harvested and mixed in a ratio of 70/30 (vol/vol) with the conditioned medium from the nerve terminal chamber of another microfluidic device seeded with naïve hippocampal neurons. This mix of media was consequently applied either to the soma chamber or nerve terminal chamber of the device containing naïve hippocampal neurons. On DIV 13-15, treated microfluidic devices were stimulated with a “pulse” corresponding to addition of high potassium (HK^+^) buffer (56 mM KCl, 0.5 mM ascorbic acid, 0.1% BSA, 15 mM HEPES, 5.6 mM D-Glucose, 95 mM NaCl, 0.5 mM MgCl2, 2.2 mM CaCl2, pH 7.4 and osmolarity 290-310) to the soma chamber, and addition of HK^+^ buffer containing Cholera toxin subunit B conjugated to either Alexa Fluor™ 555 or Alexa Fluor™ 647 (CTB-Af555 or CTB-Af647) (50 ng/ml) (C34776 and C34778 respectively, Invitrogen) to the nerve terminal chamber of the microfluidic device. The device was then incubated 5 minutes at 37°C before a washing step consisting of three washes with low potassium (LK^+^) buffer (5.6 mM KCl, 0.5 mM ascorbic acid, 0.1% BSA, 15 mM HEPES, 5.6 mM D-Glucose, 145 mM NaCl, 0.5 mM MgCl2, 2.2 mM CaCl2, pH 7.4 and osmolarity 290-310) to remove excess of CTB that has not been endocytosed. Finally, the LK^+^ buffer was replaced by the original conditioned medium and the device was put back at 37°C for a 2h chase. After the chase, live imaging of CTB-Af555 or CTB-Af647 undergoing retrograde transport in axons crossing the grooves of the microfluidic device was performed using a microscope equipped with an incubation system allowing to perform imaging at 37°C and 5% CO2 (Diskovery spinning disk confocal microscope or a Zeiss Elyra SIM/PALM/STORM super-resolution microscope).

### Depletion of α-syn from conditioned medium

PC12 cells were co-transfected with either α-syn^WT^-GFP and Hsp90^WT-^mCherry or α-syn^WT^-GFP and Hsp90^MD7-^mCherry. Following 24h transfection, conditioned medium from cells expressing α-syn^WT^-GFP and Hsp90^WT-^mCherry was collected and immunodepleting was conducted using GFP-Trap^®^ Magnetic Agarose beads (Chromotek) according to manufacturer’s protocol. Briefly, beads were incubated with the conditioned medium (25 µL of beads for 1 mL of medium) for 1 hour at 4°C under rotation. Magnetic beads were then separated by placing tubes containing the conditioned medium on a magnetic rack and removing the supernatant. Afterwards, the supernatant depleted of free α-syn was incubated with naïve neurons seeded in microfluidic devices to analyse the effect of the depleted medium on retrograde transport of CTB in the recipient neurons (see section “Neuronal treatment, stimulation, and nerve terminal labelling with CTB” for detailed protocol). Imaging was performed on a Diskovery spinning disk (see section “Diskovery spinning disk confocal imaging and analysis” for details). Immunodepleted conditioned medium was assessed for α-syn by running on SDS-PAGE gel and Western-blotting. In brief, conditioned medium samples were solubilised in either 2X sodium dodecyl sulfate (SDS) with 10% β-mercaptoethanol (for the original medium) or in 1X SDS with 10% β-mercaptoethanol (for the depleted medium) and boiled at 95 °C for 5 minutes. Equal amounts of proteins were then loaded on a Tris-Glycine precast gel (4–15%, Bio-Rad), ran at 100 V for an hour and subsequently transferred onto a Immobilon^®^-FL PVDF membrane (05317, Millipore) and incubated one hour in blocking buffer (Odyssey^®^ Tris-Buffered Saline, Li-Cor). Afterwards, membranes were incubated in a mix of 1X PBS/0.2% Tween^®^ 20 (P1379, Sigma-Aldrich) with primary antibodies overnight at 4°C (mouse anti-α-syn, 1:250, Abcam ab27766 and rabbit anti-GFP, 1:1000, Millipore AB3080P). Membranes were then washed 3 times for 10 minutes with Tris-Buffered Saline/0.2% Tween^®^ 20 (P1379, Sigma-Aldrich) and incubated in blocking buffer (Odyssey^®^ Tris-Buffered Saline, Li-Cor) containing InfraRed secondary antibody diluted 1:20,000 (goat anti-rabbit IRDye^®^ 680LT, 926-68021 and goat anti-mouse IRDye^®^ 800CW, 926-32210, Millennium Sciences).

### Diskovery spinning disk confocal imaging and analysis

Time-lapse imaging was performed at the Queensland Brain Institute’s Advanced Microscopy Facility using a Nikon Plan CFI Apo Lambda 60x Oil objective (Numerical Aperture 1.45, W.D. 0.13mm) on a spinning disk confocal microscope (Diskovery; Andor Technology, UK), equipped with 4 laser lines (405nm, 588nm, 561nm and 640nm), 500µm wide range piezo z-drive, two Zyla 4.2 sCMOS cameras (Andor Technology), and built around a Nikon Ti-E body (Nikon Corporation, Japan) and controlled by Nikon NIS software (version 5.2). Live movies were acquired at 3.3 Hz with 300 ms exposure using the 100nm pinhole. Movies were then deconvoluted using Huygens (version 17.10-64 and 18.04-64) and analysed using ImageJ (version 1.53) with the Trackmate plugin (version 3.4.2) to track the number of CTB-positive carriers. The data generated by Trackmate were then processed using R (1.2.1335) to obtain final data on the frequency of CTB+ carriers. Kymographs were generated using the KymographBuilder plugin (version 2.1.1) on ImageJ.

### Exosomal release live imaging and analysis

48h after transfection, PC12 cells or hippocampal neurons (DIV 18) cultured in glass-bottom dishes were washed once with imaging buffer (145 mM NaCl, 5 mM KCl, 1.2 mM Na2HPO4, 20 mM HEPES and 10 mM D-glucose, pH 7.4) or LK+ buffer (5.6 mM KCl, 0.5 mM ascorbic acid, 0.1% BSA, 15 mM HEPES, 5.6 mM D-Glucose, 145 mM NaCl, 0.5 mM MgCl2, 2.2 mM CaCl2, pH 7.4 and osmolarity 290-310) for PC12 cells or neurons respectively, before imaging at the Queensland Brain Institute’s Advanced Microscopy Facility using the Roper iLas 2 microscope (generously supported by the Australian Government through the ARC LIEF grant LE130100078). For experiments characterising the effect of Hsp90^MD7^ on exosomal release in PC12 cells, prior to each time-lapse recording, Hsp90^MD7^expression level in the cell was verified by acquiring an image in the red (561) channel. Analysis of exosomal release was performed using ImageJ (version 1.53). Where needed, raw movies were drift corrected using the Stabilizer plugin. Exosomal release in PC12 cells transfected with CD63-pHluorin was characterised by a burst in fluorescence in the green (488) channel, corresponding to the fusion of MVB with the plasma membrane leading to neutralisation of the pH in the MVB lumen and unquenching of fluorescence. One burst of fluorescence was called a “fusion event”. For each recording, the number of fusion events and the duration of each event were determined. The duration of an event was defined from the time point fluorescence started to increase until the fluorescence had returned to the basal level. When there was no return to initial fluorescence, the end time point was determined as the time when the fluorescence reached a steady state. Regarding exosomal release of α-syn in hippocampal neurons, analysis of exocytic events was performed with the Bar plugin of ImageJ, which provided measurements of fluorescence over time in both red and green channels in hippocampal neurons transfected with CD63-pHuji and α-syn-GFP. A “fusion event” was defined by a sharp decrease in fluorescence in the red channel. For each recording, the number of fusion events and their dynamics (event duration, rise time, decay time, maximum fluorescence, amplitude of fluorescence variation) were determined. The duration of an event was defined in the red channel from the time point fluorescence started to decrease until the moment fluorescence had reached a plateau. In the green channel, an event was defined from the time point fluorescence started to increase until the time fluorescence had returned to the basal level or reached steady state if there was no decrease.

### TIRF microscopy – Roper iLas 2 Microscope

Time-lapse TIRF (total internal reflection fluorescence) imaging was performed on a motorized inverted microscope (Nikon, TiE) equipped with a perfect focus system with an ILas2 double-laser illuminator (Roper Scientific), two Evolve 512 delta EMCCD cameras (Photometrics) mounted on a TwinCam LS Image Splitter (Cairn Research) and an incubation system. The microscope has 2 sets of optical filters, one being within the microscope’s body: a QUAD dichroic filter (ZT405/488/561/647rpc, Chroma) with a QUAD emission filter (ZET405/488/561/640m, Chroma), the second set being within the splitter. The splitter contains a dichroic filter and two emissions filters which can be changed depending on the experimental design. The classic setting comprises the dichroic ZT647rdc-UF1 (Chroma) and two emission filters (Chroma): ET690/50m and ZET405/488/561m. For dual live imaging of α-syn-GFP and CD63-pHuji, the dichroic filter and emission filters on the splitter were changed for: the dichroic filter T565lpxr (Chroma), emission filters ET600/50m and ET540/40m (Chroma). Prior to imaging, alignment of cameras and TIRF angle calibration were done using TetraSpeck™ microspheres beads (T729, Invitrogen). Neurons were maintained at 37°C and 5% CO_2_ during imaging. Time-lapse movies were acquired using a TIRF angle allowing selective imaging of the plasma membrane. Laser power was set at 6.5% for the 561 channel and 10.5% for the 488 channel and recording was performed at 2 Hz with 20 ms exposure over 10 minutes. Stimulation was performed at 5 minutes by adding either 2mM barium chloride (100474N, BDH Reagents & Chemicals) to PC12 cells, or HK+ buffer (56 mM KCl, 0.5 mM ascorbic acid, 0.1% BSA, 15 mM HEPES, 5.6 mM D-Glucose, 95 mM NaCl, 0.5 mM MgCl2, 2.2 mM CaCl2, pH 7.4 and osmolarity 290-310) to hippocampal neurons. Image acquisition was performed using Metamorph (version 7.10.1.161; Molecular Devices).

### Single-particle tracking photoactivated localization microscopy (sptPALM)

A day after transfecting hippocampal neurons (DIV 11-13) of the soma chamber of microfluidic devices with either mEos2 alone, α-syn^WT^ -mEos2 or α-syn^A30P^-mEos2, conditioned media from the soma chambers were collected and mixed with conditioned media from the nerve terminal chamber of naïve recipient neurons (70/30 ratio) before adding the mix to recipient neurons of the nerve terminal chamber (DIV 12-14) for 24 hours. After a 5-minute pulse and 2h chase, time-lapse imaging was performed at the Queensland Brain Institute’s Advanced Microscopy Facility using an Alpha Plan-Apochromat 100X (1.46 NA) oil objective on the Zeiss ELYRA SIM/PALM/STORM microscope equipped with an Andor 897 EMCCD camera and an incubation system allowing imaging under 37°C and 5% CO_2_ conditions. Time-lapse movies were acquired using a TIRF (total internal reflection fluorescence) angle allowing selective imaging of the plasma membrane. Axons were located using the 642 nm laser to visualise CTB-Af647 uptaken in neurons. Imaging of CTB was done with 1% laser power, 300 ms exposure time over 400 frames. Subsequently, imaging of transmitted mEos2 molecules within neurons was achieved using the 405 nm and 561 nm lasers to respectively photoconvert mEos2 molecules and visualise them. The 405 nm laser was used at 0.0002-0.3% power, whereas the 561 nm laser was used at 30% power. Recording was performed over 2000 or 16000 frames and 30 ms exposure time. Movie acquisition was done using the Zen acquisition software (Black edition, 2012 version, Carl Zeiss).

### sptPALM analysis and cluster analysis

ImageJ (version 1.53) was used to convert movies from Zen’s CZI format to a TIFF format. The localisation and mobility of single molecules was determined using a custom written PALMtracer plugin in Metamorph (version 7.10.1.161; Molecular Devices) (Kechkar et al. 2013) (Nair et al. 2013). The analysis was done for specific regions of interest (ROI) as well as for the whole field of view. Regions of interest were drawn around axons based on the maximum Z projection image of CTB-Af647 molecules moving within axonal projections situated in the nerve terminal chamber of microfluidic devices. PALM-Tracer generated trajectory maps, super-resolved average intensity, and diffusion coefficient (D) maps as well as excel files containing diffusion coefficient and mean square displacement (MSD) values. Thresholding of the signal was done individually for each movie. Only molecules picked up by the thresholding and which fluorescence lasted across eight frames of the movie were tracked. The mean square displacement was determined for each trajectory using the equation *MSD* (*t*) = *a* + 4*Dt* where “a” is the offset constant which incorporates the effects of localization error and finite camera exposure, “D” is the diffusion coefficient and “t” is the time. The first eight points of the MSD were averaged for all trajectories and plotted against time. The area under the MSD curve (AUC) was used to quantify the significancy of changes in MSD values. Molecules were distributed in mobile and immobile populations based on the value of the diffusion coefficient where log10(D)> −1.5 µm^2^s^−1^ was considered the mobile fraction (Constals et al. 2015). The percentage of uptake of molecules within neurons was determined using the following formula: ((Number of tracks. µm^−1^ in neurons) - (Number of tracks.µm^−1^ without ROI) / (Number of tracks.µm^−1^ in neurons))*100. The periodicity analysis of clusters was done on the intensity average map using ImageJ where a line was drawn between each cluster or from one edge to the other of a cluster to respectively determine the spacing between clusters or the cluster size. In addition, based on data generated from PALM-tracer, a cluster analysis was performed using the Nanoscale Spatiotemporal Indexing Clustering (NASTIC) tool as previously described (Wallis et al. 2021).

### Plasmids and Fluorescent markers

EGFP-C1 and mCherry-N1 (632523) were purchased from Clontech and mEos2-N1 was purchased from Addgene (54662). α-syn^WT^-GFP and α-syn^A30P^-GFP (GFP on the N-terminus) were previously developed in the laboratory by Dr. Ye Jin Chai (Chai et al. 2016). Hsp90^WT^-mCherry and Hsp90^MD7^-mCherry (mCherry on the C-terminus) were kindly provided by Dr. Patrik Verstreken (VIB-KU Leuven Center for Brain & Disease Research, Belgium). Hsp90^MD7^ has three mutations in its amphipathic helix: K392A, K395A and K399A. CD63-pHluorin and CD63-pHuji plasmids were kindly provided by Dr. Dirk Michiel Pegtel (Cancer Center Amsterdam, VU University Medical Center, Netherlands).

### Reagents

The following reagents were sourced at Sigma-Aldrich: NaCl (746398), KCl (P3911), MgCl_2_ (M9272), CaCl_2_ (C5080), HEPES (H3375) ascorbic acid (A7506), Bovine serum albumin (BSA) (A8022). D-glucose was purchased from Amresco (0188). Anti-GFP magnetic beads were purchased from ChromoTek (GFP-Trap^®^ Magnetic Agarose beads, gtma-20).

### Statistical analysis

GraphPad Prism 9.4.1 (San Diego, USA) was used to determine statistical significance and determine outliers using ROUT method. Normal distribution of data was tested using the “D’Agostino-Pearson omnibus” normality test. For comparison between two groups that were normally distributed, a student t test was used. For unpaired non-parametric data sets, a Mann-Whitney test was used, or Wilcoxon signed-rank test was used for paired data sets. For datasets which compared more than 2 groups, an ANOVA was used with multiple comparisons (for normally distributed data, an ordinary one-way ANOVA with Tukey’s multiple comparisons was performed; for non-parametric data, Kruskal-Wallis test with Dunn’s multiple comparisons was performed). Tests used are indicated in the respective figure legends. P values < 0.05 were considered statistically significant.

## Data Availability

All processed data will be made available upon request and into The University of Queensland data repository, UQ eSpace.

## Authors’ contributions

All authors contributed to the study by performing experiments, data analysis, interpretation, by supplying reagents or writing the manuscript. F.A.M conceived and B.E.D and V.L coordinated and performed the study. B.E.D prepared the figures and wrote the manuscript with F.A.M, R.S.G, T.P.W and A.J.M with editing and input from co-authors. V.L performed the immunohistochemistry, the immunodepletion of the conditioned medium and Western Blot as well as all CTB retrograde transport experiments and their analyses. B.E.D performed Hsp90 characterisation experiments, the exosomal release experiments as well as the super resolution experiments and their analyses. A.T.B. provided technical support for the super-resolution experiments. R.S.G, T.P.W and A.J.M gave input on figures and participated in manuscript writing. J.J.C-W provided the microfluidic devices. P.V provided the Hsp90 constructs.

## Acknowledgments

The authors acknowledge the use of the Australian Microscopy & Microanalysis Research Facility at The University of Queensland. We thank IT department and the Advanced Microscopy and Microanalysis Facility at the Queensland Brain Institute for their outstanding support. We thank Dr. Dirk Michiel Pegtel for providing the CD63 constructs. The work was supported by an Australian Research Council (ARC) Discovery Project grant (DP170100125), an ARC Linkage Infrastructure Equipment and Facilities grant (LE130100078) and a National Health and Medical Research Council (NHMRC) Senior Research Fellowship (1155794) to F.A.M. The Research Training Program (RTP) Scholarship and a QBI top-up Scholarship supported B.E.D work.

---

**Supplementary Figure 1:**
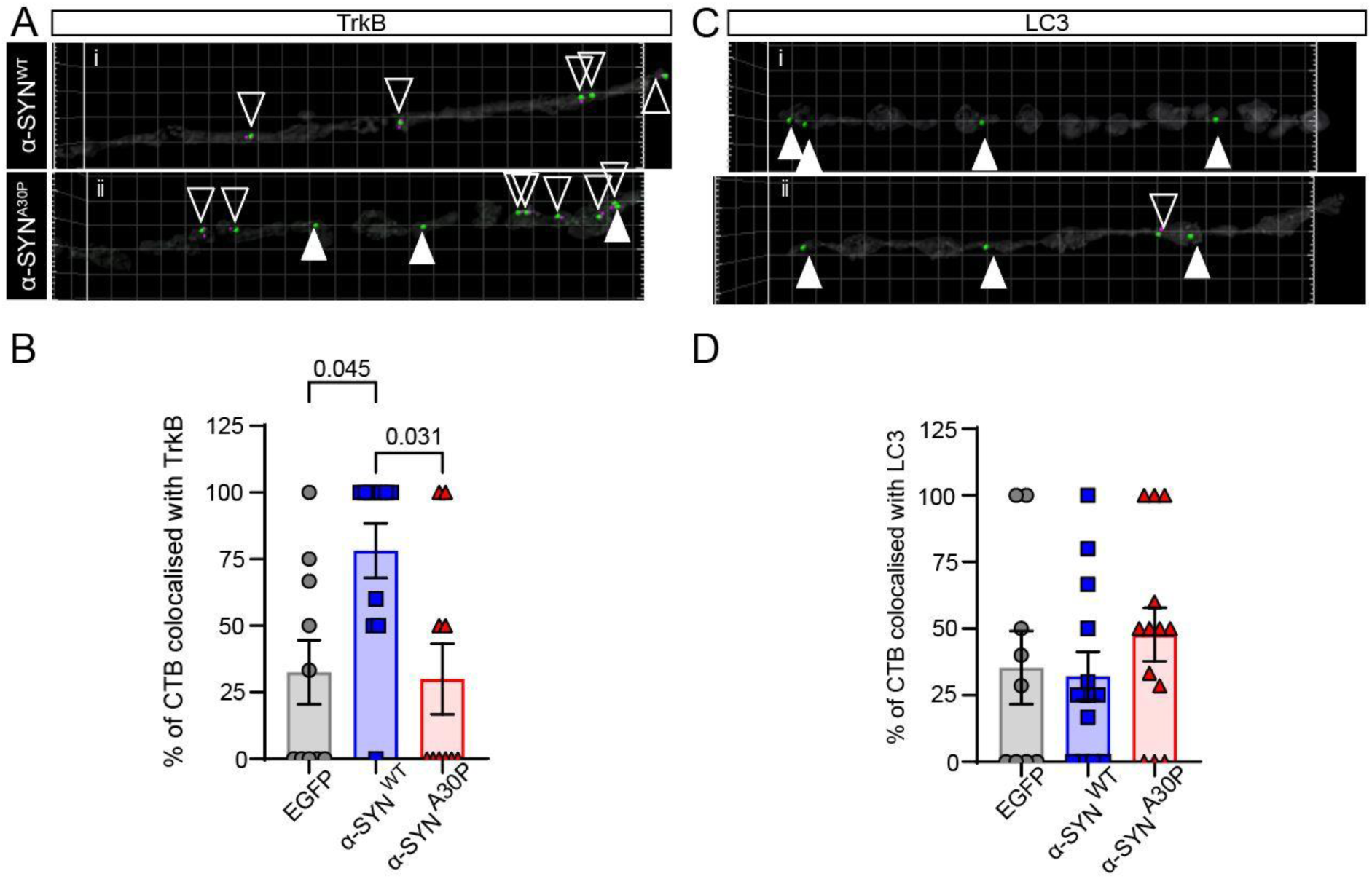
Conditioned medium from hippocampal neurons expressing α-syn^WT^-GFP increases colocalization of axonal CTB with TrkB in recipient neurons. **A**. Representative image showing the axonal colocalization of CTB-Alexa647 with TrKB (green) in hippocampal neurons preincubated with α-syn^WT^-GFP (Top) or α-syn^A30P^-GFP (bottom). **B**. Quantification of CTB-positive carriers colocalizing with TrkB in hippocampal neurons preincubated with α-syn^WT^-GFP or α-syn^A30P^-GFP. **C**. Representative image showing the axonal colocalization of CTB-Alexa647 with LC3 (green) in hippocampal neurons preincubated with α-syn^WT^-GFP (Top) or α-syn^A30P^-GFP (bottom). **D**. Quantification of CTB-positive carriers colocalizing with LC3 in hippocampal neurons preincubated with α-syn^WT^-GFP or α-syn^A30P^-GFP. Data are from 2 independent experiments, with 13–18 axonal channels imaged per condition. Results are presented as mean ± SEM and analyzed by one-way ANOVA with Tukey’s post hoc test.

**Supplementary Figure 2:**
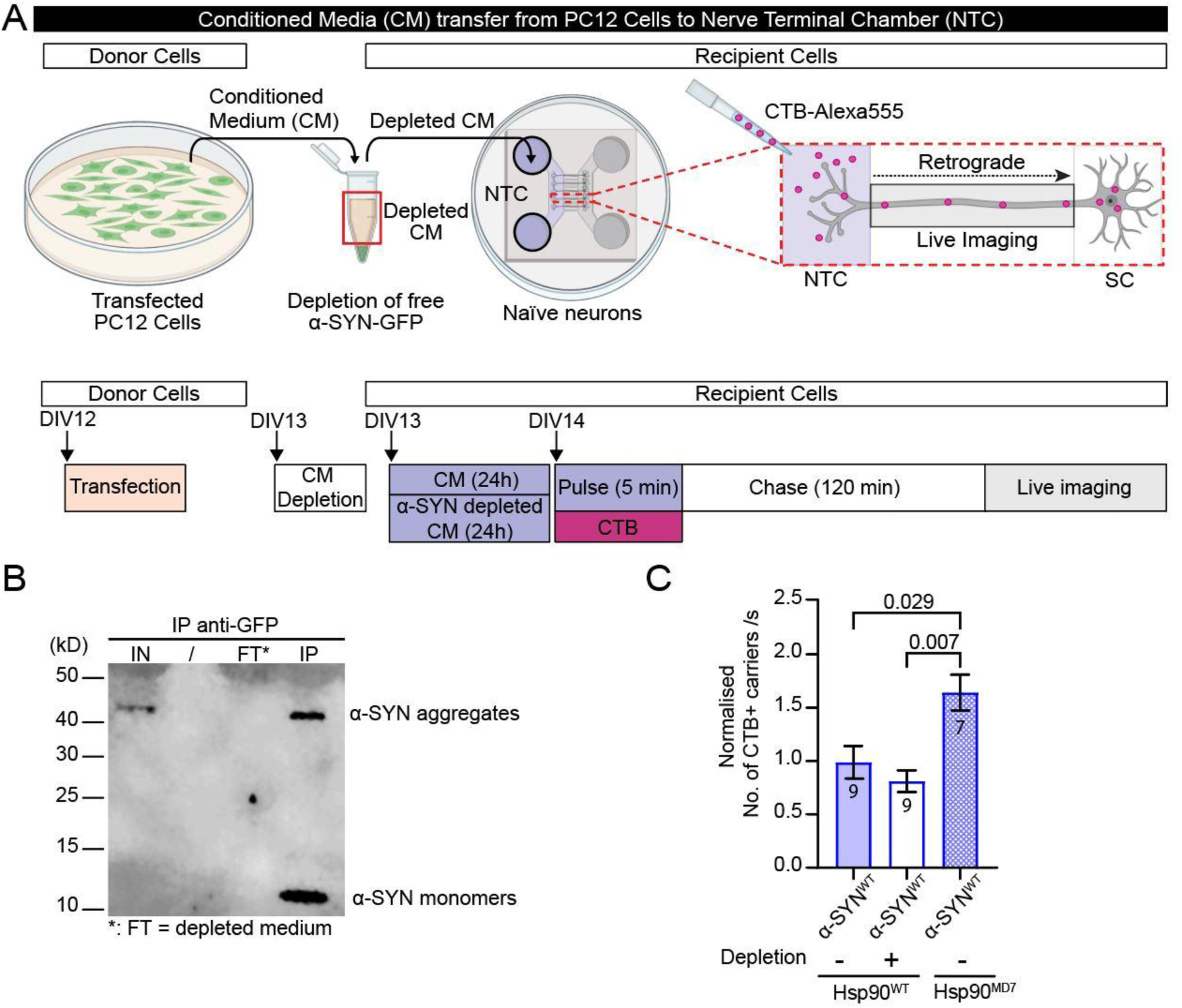
The downregulating effect of the conditioned medium in recipient cells is not mediated by free secreted α-syn. **A.** Schematic of the experimental paradigm: PC12 cells were co-transfected with either α-syn^WT^-GFP and Hsp90^WT^-mCherry or α-syn^WT^-GFP and Hsp90^MD7^-mCherry (donor device; DIV 12). After 24h, part of the conditioned medium from PC12 cells co-transfected with α-syn^WT^-GFP and Hsp90^WT^-mCherry was immunodepleted of α-syn-GFP using anti-GFP magnetic beads. Subsequently, each conditioned medium (α-syn^WT^-GFP/Hsp90^WT^ original and immunodepleted samples as well as was α-syn^WT^-GFP/Hsp90^MD7^) was added to the nerve terminal chamber (NTC) (70:30 ratio) of a microfluidic device containing untransfected naïve neurons. The following day a 5-minute pulse of cholera toxin B subunit Alexa Fluor™ 555 (CTB-555; 50 ng/mL) diluted in high potassium (high K^+^) buffer was conducted followed by 3 washes with low potassium (LK^+^) buffer to remove the excess of CTB. Afterwards, devices were incubated back with the conditioned media for 2h before imaging CTB-positive carriers undergoing retrograde transport in axons of the recipient neurons. **B.** Western Blot of the conditioned media from PC12 cells co-transfected with α-syn^WT^-GFP and Hsp90^WT^-mCherry before and after immunodepletion to confirm the efficient removal of α-syn. IN = input corresponding to the loading of the original conditioned medium, FT = flow-through corresponding to the loading of α-syn depleted fraction, IP = immunoprecipitation corresponding to the loading of the fraction containing α-syn-GFP bound to magnetic beads. **C.** Quantification of the frequency of CTB+ carriers (number of carriers per second). Timelapse confocal imaging was performed on the Discovery spinning disk at a 3Hz imaging rate. Data from 1 independent experiment, n=7-9 channels, Kruskal-Wallis test followed by Dunn’s comparison test, mean ± SEM.

**Supplementary Figure 3:**
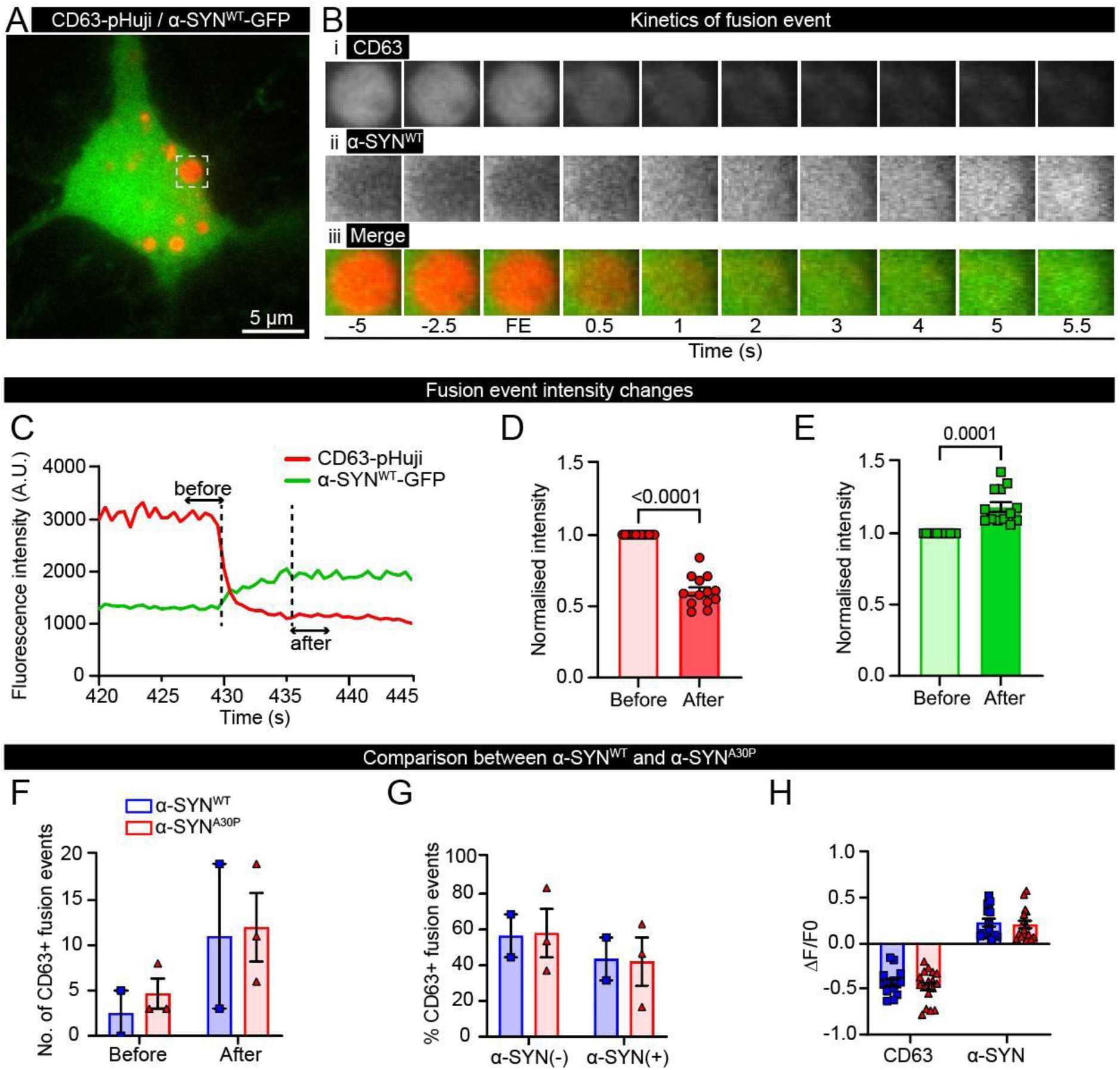
Live imaging of α-syn exosomal release in hippocampal neurons does not reveal any differences between WT and A30P mutant. Scheme of the experiment: Glass-bottom dishes seeded with hippocampal neurons extracted from brains of E16 mice embryos were co-transfected with exosomal marker CD63-pHuji and α-syn^WT^-GFP at DIV 18. At DIV 20, neurons were washed in low potassium buffer, and TIRF live-cell dual imaging was performed for 10 min with stimulation after 5 minutes using high potassium buffer. CD63-pHuji is quenched in the acidic environment of the multivesicular body (MVB). Upon fusion of MVB with the plasma membrane, CD63-pHuji becomes exposed to the neutral extracellular environment and is unquenched, enabling visualisation of CD63-pHuji proximal to the membrane via TIRF. Then, exosomal α-syn-GFP is released, reflected by a sharp decrease in the red channel fluorescence (CD63-pHuji) and an increase in the green channel (α-syn-GFP). **A.** TIRF fluorescent image of a hippocampal neuron expressing α-syn-GFP (green) and CD63-pHuji (red) with one MVB outlined with dashed line, scale bar: 5 µm. **B-E.** Characterisation of the fusion with the plasma membrane of the MVB outlined in (A). **B.** Timelapse cartoon of the exocytic event, time in seconds. i. Representative images of fluorescence intensity changes in the red channel (CD63-pHuji); ii. Representative images of fluorescence intensity changes in the green channel (α-syn-GFP); iii. Representative images of fluorescence intensity changes in red and green channels. **C.** Graph illustrating the method used to characterise changes in fluorescence during a fusion event. For each fusion event, the mean fluorescent intensity prior to the beginning of the fusion event was determined over 10 frames, as well as the mean intensity over 10 frames at the end of the fusion event, corresponding to the maximum value in the green channel (α-syn-GFP), and the minimum value in the red channel (CD63-pHuji). **D.** Graph of normalised intensities in the red channel before and after the exocytic event. Fluorescent intensities were normalised to the intensity at the beginning of the exocytic event. **E.** Same as D. but in the green channel. **F.** Graph comparing the number of fusion events in the red channel before and after stimulation. **G.** Graph representing the percentage of exocytic events that contained α-syn and ones that did not. **H.** Graph representing amplitude of changes in fluorescence before and after each exocytic event. Values were normalised to the intensity measured at the beginning of an exocytic event.

